# The *C. elegans* piRNA pathway safeguards splicing and RNA fidelity

**DOI:** 10.1101/2025.10.27.684940

**Authors:** Sehee Min, Johnny Cruz-Corchado, Veena Prahlad

## Abstract

Piwi-interacting RNAs (piRNAs) silence transposons and foreign sequences; yet, for poorly understood reasons, they also extensively target endogenous mRNAs. Here, we demonstrate that this targeting preserves transcript isoform fidelity by regulating pre-mRNA splicing in *C. elegans*. Using the heat shock response to provoke rapid transcriptional changes, we found that piRNAs and PIWI/PRG-1 direct the production of endogenous antisense 22G-RNAs from newly induced transcripts. These endo-siRNAs associate with RNA Polymerase II, spliceosome protein MOG-7/C2orf3/Ntr2, and nascent RNAs to delay splice completion and regulate splice-site selection. Long-read transcriptome sequencing revealed that loss of piRNAs accelerates splice completion but compromises accurate splice-site selection, increasing the production of aberrant transcript isoforms that are normally eliminated by alternative splicing coupled to nonsense-mediated decay (AS-NMD). Consequently, piRNA pathway mutants are profoundly reliant on NMD for embryonic survival, development, and proteostasis maintenance. Together, our findings identify a previously unrecognized role for the piRNA pathway in coordinating splicing with RNA quality control to preserve proteome integrity. We propose that this unexpected role for the piRNA pathway may have evolved from its ancestral function in recognizing transposable elements, which frequently introduce cryptic splice sites and undergo exonization, to generate novel transcripts that require elimination by NMD.

## Introduction

Accurate pre-mRNA splicing is essential for maintaining transcript identity and proteome integrity^1–4^. Yet, how cells ensure splice-site fidelity and coordinate splicing kinetics with changing physiological demands—particularly during stress and rapid transcriptional remodeling, when RNA processing capacity can become limiting^5–10^— remains poorly understood. Here, we report a previously unrecognized role for PIWI-interacting RNAs (piRNAs) in this process.

piRNAs are small, single-stranded RNA guides for PIWI Argonaute proteins, best known for protecting genomes by suppressing transposable elements and foreign nucleic acids^11–19^. Yet, across animal species, for reasons that remain largely unknown, a large fraction of piRNAs also target endogenous mRNAs^11–20^. One well-established consequence of this targeting is transcriptional and post-transcriptional gene silencing^11,12,21–23^. In *C. elegans*, 15,000–17,000 distinct piRNA (21U-RNAs) are constitutively expressed from two genomic regions on Chromosome IV, called piRNA clusters^24–27^. These piRNAs associate with PRG-1, the sole *C. elegans* PIWI protein^26^, and owing to their high mismatch tolerance, can target nearly all expressed RNAs — including transcripts never previously encountered, such as those from GFP transgenes^11,12,26,28–31^. The piRNA-targeting event is amplified through the synthesis of abundant secondary antisense small endogenous silencing RNAs (endo-siRNAs, or 22G-RNAs) generated by RNA-dependent RNA polymerases (RdRP), which use the piRNA-targeted mRNAs as templates for antisense 22G-RNA synthesis. These 22G-RNAs then serve as guides for other Argonaute proteins^21,29,30^. Consequently, *C. elegans* constitutively produces an extraordinarily complex population of 22G-RNAs that collectively survey much of the transcriptome. Yet, most mRNA targets of piRNAs that generate 22G-RNAs are neither foreign nor ultimately silenced, raising the fundamental question of why the piRNA pathway monitors such a large fraction of endogenous transcripts.

To address this question, we exploited the heat shock response to provoke widespread transcriptional changes^5–7^— inducing the massive expression of previously undetectable heat-shock genes (*hsps*) while altering the levels of almost all other existing transcripts— and asked how the piRNA system responds to this pervasive transcriptome remodeling. We found that heat shock stimulated the expression of select piRNAs and the production of piRNA-dependent 22G-RNAs from more than 40% of induced genes, including the *hsp* genes themselves. Unexpectedly, rather than promoting post-transcriptional silencing of the induced mRNAs, these newly generated piRNA-dependent antisense 22G-RNAs associated with RNA polymerase II (Pol II), nascent transcripts, and the spliceosomal protein MOG-7, the *C. elegans* functional ortholog of Ntr2/C2ORF3, to delay splice completion and promote accurate splice-site selection. Consistent with a role in splicing, long-read transcriptome sequencing revealed that loss of piRNAs led to widespread alterations in transcript architecture, exon usage, and splice-site choice, increasing the production of aberrant transcript isoforms that are normally eliminated by alternative splicing coupled to nonsense-mediated decay (AS-NMD). Consequently, loss of the piRNA pathway rendered animals profoundly reliant on NMD, and the genetic disruption of nonsense-mediated decay (NMD) in piRNA-deficient animals led to the accumulation of misfolded proteins and synthetic embryonic lethality. Finally, we found that this regulatory pathway appeared to be integrated into the heat-shock response: the conserved stress-responsive transcription factor HSF-1 localizes to piRNA clusters, promotes expression of piRNAs predicted to target *hsp* transcripts, coupling transcriptional activation to alternative splicing and RNA quality control during proteotoxic stress.

Together, our findings identify a previously unrecognized role for the *C. elegans* piRNA pathway in regulating splicing fidelity and, thereby, maintaining proteome integrity, suggesting that this might provide an explanation for the piRNAs’ surveillance of endogenous genes. We speculate that this function may have evolved from the ancestral ability of piRNAs to recognize transposable element-derived sequences that are frequently embedded within introns, and when used, generate alternatively spliced isoforms eliminated by NMD^32–36^.

## Results

### Heat-shock-induced *hsp* and other mRNAs are targeted by the germline piRNA pathway

To determine how the piRNA surveillance system responds to pervasive changes in the mRNA repertoire of cells, we performed mRNA (paired-end, strand-specific, polyA-selected) and small RNA sequencing (sRNA-seq) on synchronized day-one adults exposed to short (5 min, 34°C) or prolonged (30 min, 34°C) heat shock, followed by a 2-hour recovery period when normal physiological functions largely resume^37,38^ (Supplementary Fig. 1a; Supplementary Tables 1 and 2). To distinguish piRNA-dependent effects, we compared wild-type animals with *prde-1* mutants, which are defective in piRNA biogenesi.

As expected, in wild-type animals, heat shock triggered extensive transcriptional changes, resulting in the differential expression (p. adj < 0.05) of ∼3000 mRNAs by 30 minutes of heat shock exposure (Fig. 1a; Supplementary Fig. 1b, c). Unexpectedly, nearly half of these heat-induced differentially expressed genes (DEGs) had generated antisense 22G-RNAs (Fig. 1a; Supplementary Table 3). This included the newly induced *hsp* genes (Fig. 1b; Supplementary Fig. 1d). In wild-type animals, 22G-RNAs antisense to *hsps* were produced within 5 minutes of heat shock, increased in abundance with 30 minutes of heat shock, and further increased during recovery from heat shock. Their kinetics mirrored the accumulation of mature *hsp* mRNAs, which also peaked ∼2–4 hrs post-heat shock during recovery (Supplementary Fig. 1e), even though *hsp* transcription occurred almost entirely during the 30-minute heat-shock exposure, as evidenced by Pol II occupancy at the *hsp* genes, *hsp-70s* (*C12C8*.1 and *F44E5*.4/.5; ChIP-qPCR; Supplementary Fig. 1f). Consistent with 22G-RNA biogenesis requiring mRNAs as templates for RdRP activity, a mutation in HSF-1, *hsf-1 (sy441)*, that impairs heat-shock-induced mRNA expression also reduced heat-induced 22G-RNA production (Fig. 1a; Supplementary Fig. 1c, d), including those templated by *hsp* mRNAs (Fig. 1b; Supplementary Fig. 1d). Notably, even though *hsp-70 (F44E5.4/.5)* mRNA was induced in *hsf-1* mutants, it failed to produce 22G-RNAs, suggestive of a broader role for HSF-1 in this response (Supplementary Fig. 1d). Together, these findings indicated that the heat-shock-induced, HSF-1-dependent, newly synthesized transcripts, including *hsps*, served as templates for antisense 22G-RNA generation and might engage the piRNA surveillance system.

**Figure 1.**
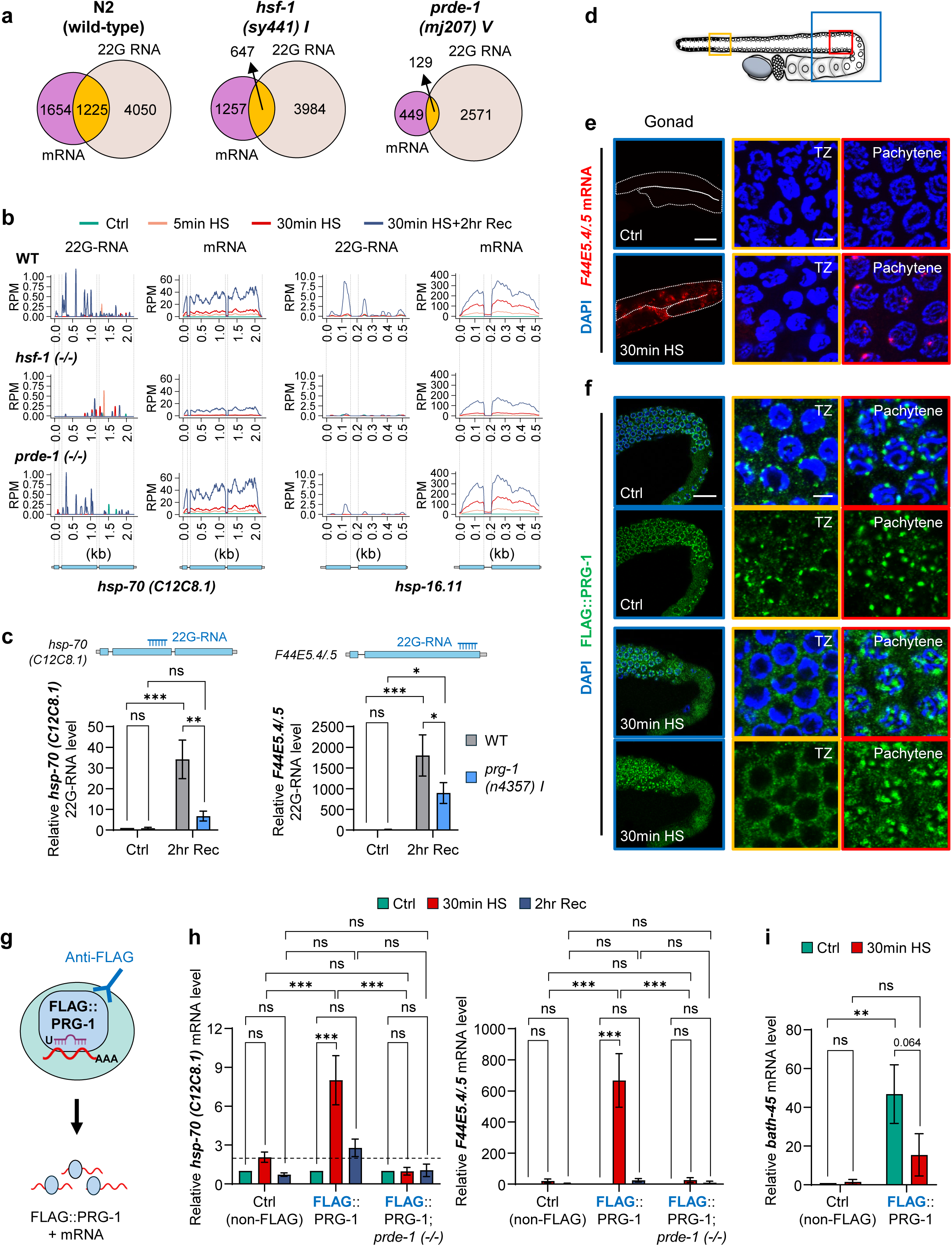
Heat-shock-induced *hsp* mRNAs are targeted by the germline Piwi-interacting RNAs (piRNAs) pathway. **a.** Overlap between heat-shock-induced mRNAs and antisense 22G-RNAs in wild-type (N2), *hsf-1* (*sy441*), and *prde-1* (*mj207*) animals. Numbers indicate significantly induced mRNAs and 22G-RNAs and their overlap. (30min HS). **b.** 22G-RNA and mRNA profiles across *hsp-70* (*C12C8.1*) and *hsp-16.11* in wild-type, *hsf-1*, and *prde-1* animals under control conditions, after 5 or 30 min heat shock (HS), and after 30 min HS followed by 2 hr recovery (Rec) are shown below. **c.** Stem-loop RT–qPCR quantification of 22G-RNAs antisense to *hsp-70 (C12C8.1)* and *F44E5.4/.5* in wild-type and *prg-1 (n4357)* animals under control conditions and after HS followed by 2 hr recovery. *n* = 4 biological repeats. **d–f.** Germline localization of *hsp-70* transcripts and PRG-1. **(d)** Schematic showing the gonadal regions imaged. **(e)** Representative projected confocal images of RNA Fluorescence *in situ* hybridization (RNA FISH) for *F44E5.4/.5* mRNA (red) before and after 30 min HS. **(f)** Representative confocal Z-sections FLAG::PRG-1 (green) before and after 30 min HS. Enlargements show the transition zone (TZ) and pachytene regions. DNA, DAPI (blue). Scale bars, 50 μm (Gonad; e), 20 μm (Gonad; f), and 3 μm (TZ, Pachytene). **g–i.** PRG-1 association with heat-shock-induced transcripts. **(g)** Schematic of FLAG::PRG-1 RNA immunoprecipitation (RIP). **(h)** Fold changes in *hsp-70s* (*C12C8.1* and *F44E5.4/.5*) mRNA level co-immunoprecipitated with PIWI/PRG-1 in wild-type and *prde-1* mutant animals under control, 30 min HS, and 2 hr Rec conditions, relative to the control conditions of each strain. **(i)** *bath-45* mRNA co-immunoprecipitated with PRG-1 before and after HS, relative to the control condition of non-FLAG controls. mRNA levels were normalized to *pmp-*3 (input). *n* = 3–11 biological repeats. **c, h, i**: Data are mean ± SEM. Statistical analyses: two-way ANOVA. ns, not significant; ✱ *p* < 0.05; ✱✱ *p* < 0.01; ✱✱✱ *p* < 0.001.

Examination of *prde-1* mutants confirmed that the 22G-RNAs generated from HSF-1-induced transcripts were piRNA-dependent. Although *prde-1* mutants, which lack piRNAs under basal and heat-shock conditions (Supplementary Fig. 2a), induced *hsp* mRNAs normally upon 30-minute heat shock, the *hsp* mRNAs produced markedly fewer 22G-RNAs (Fig. 1b; Supplementary Fig. 1d). Likewise, fewer heat-shock-responsive genes produced 22G-RNAs (Fig. 1a; Supplementary Fig. 1c). This defect was not readily attributable to a reduced germline or a global depletion of endogenous 22G-RNAs. The 1061 germline-enriched genes^39,40^ examined, were expressed at comparable levels in wild-type and *prde-1* animals (Supplementary Fig. 2b), arguing against a gross decrease in germ cell nuclei; the overall expression of DEGs, which included chaperone genes, closely resembled that in wild type (Supplementary Fig. 2b); and basal non-heat shock 22G-RNA expression did not significantly differ from wild-type except for the previously reported reduction in WAGO-associated 22G-RNAs^22,23,41,42^ (Supplementary Fig. 2c, d). Instead, the predominant defect appeared to be a selective failure to generate heat-induced 22G-RNAs (Supplementary Figs. 2e; 22G-RNAs templated by germline-expressed genes and chaperones shown). Examination of *prg-1* mutants, in which the sole *C. elegans* PIWI Argonaute^26,27^ is not functional, showed that, like *prde-1* mutants, *prg-1* animals produced markedly less heat-shock-induced 22G-RNAs antisense to *hsp-70s* (Fig. 1c), providing independent confirmation of the piRNA-pathway dependence of the heat-induced 22G-RNAs.

In *C. elegans*, canonical piRNA-dependent 22G-RNA biogenesis is thought to occur exclusively in the germline, within perinuclear germ granules (nuage), where PRG-1 and RdRPs generate 22G-RNAs from mRNAs targeted by piRNAs^26,30^. On the other hand, heat shock induces *hsp* mRNAs mainly in somatic cells, and, as has been previously shown^37,38^, in a subset of germline pachytene nuclei (Fig. 1d, e; Supplementary Fig. 2f). We therefore asked whether the heat-induced *hsp* transcripts generate 22G-RNAs in the germline through the canonical germline piRNA pathway or through other, unknown means. Immunofluorescence analyses showed that, despite its heat shock-induced dissolution from other germline regions such as the transition zone, PRG-1 remained localized in perinuclear granules in the pachytene region of the germline— the only germline region where *hsp* mRNA is also expressed upon heat shock^37,38^(Fig. 1d–f). Native stringent RNA immunoprecipitation (RIP) of FLAG-tagged PRG-1 followed by RT-qPCR demonstrated that both inducible *hsp-70* transcripts (*C12C8.1* and *F44E5.4/.5*) increased their association with PRG-1 during heat shock (Fig. 1g, h). Importantly, this was not due to non-specific binding, as levels of PRG-1-associated *bath-45*^23^, an abundant, constitutive, piRNA target, were unchanged or even decreased upon heat shock (Fig. 1i). Further, the heat-shock-induced association between *hsp* mRNAs and PRG-1 was eliminated in *prde-1* mutants that do not produce piRNAs (Fig. 1g, h). Thus, together with impaired production of *hsp*-directed 22G-RNAs in *prg-1* mutants (Fig. 1c), these findings indicate that the new stress-induced transcripts were routed through the canonical germline piRNA pathway to template piRNA-dependent 22G-RNA synthesis.

### The Heat Shock Factor 1 (HSF-1) regulates piRNA expression in response to heat shock

Because a single *C. elegans* piRNA can target multiple mRNAs^11,28^ with high mismatch tolerance, heat shock-induced 22G-RNAs could arise either from constitutively expressed piRNAs recognizing newly induced *hsp* transcripts or from the biogenesis of new piRNAs upon heat shock. To distinguish between these possibilities, we examined the expression of piRNAs predicted to target the two *hsp-70s* (Supplementary Table 4). Most detectable piRNAs predicted to target *hsp-70s* were constitutively expressed, but a small subset was also induced by heat shock in a PRDE-1-dependent manner (Supplementary Table 4). In addition, piRNAs expression was significantly increased upon heat shock (Supplementary Fig. 3a). We therefore asked whether HSF-1, the master regulator of the heat shock response, also regulates piRNA biogenesis.

Under basal conditions, piRNAs are transcribed by the PRDE-1-containing Upstream Sequence Transcription Complex (USTC) at the two chromosome IV piRNA clusters, where PRDE-1 forms condensates^24,25,43^, and HSF-1 is diffusely distributed in *C. elegans* germline nuclei (Fig. 2a). Upon heat shock, HSF-1 rapidly colocalized with PRDE-1 condensates (Fig. 2a), associating with nearly 100% of PRDE-1 condensates by 30 minutes of heat shock (Fig. 2b). In addition, HSF-1 exhibited a significant increase in chromatin occupancy at the ‘big’ piRNA cluster, where most piRNA genes reside (Fig. 2c), and upregulated 21ur-2348, predicted to target both *hsp-70* genes^11,28^ (Fig. 2d; Stem-loop RT-qPCR). In *hsf-1(sy441)* mutants, HSF-1 recruitment to PRDE-1 condensates and heat shock-induced occupancy of piRNA loci were markedly decreased (Fig. 2a), and 21ur-2348 was not induced (Fig. 2d). These findings suggested that HSF-1 might directly co-regulate stress-induced piRNA biogenesis along with PRDE-1.

**Figure 2.**
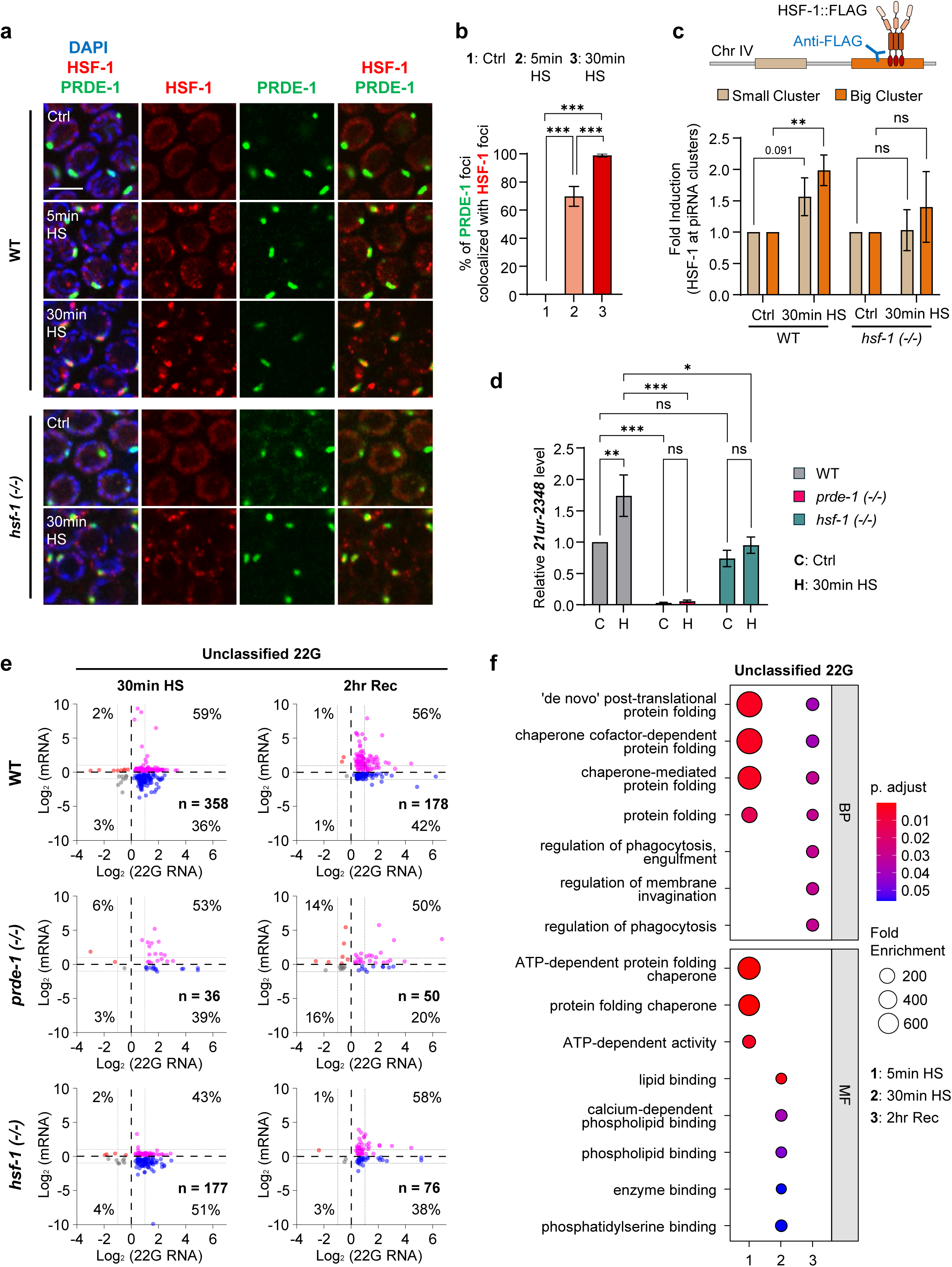
HSF-1 engages the piRNA pathway during heat shock. **a, b.** Heat shock promotes HSF-1 association with PRDE-1 nuclear foci. **(a)** Representative images of HSF-1 (red) and PRDE-1 (green) in wild-type and *hsf-1* animals under control conditions or after heat shock (HS). DNA, DAPI (blue). Scale bars, 5 μm. **(b)** Percentage of PRDE-1 foci colocalizing with HSF-1 foci after 5 or 30 min HS. *n* = 3–4 biological repeats. **c.** HSF-1 occupancy at the two chromosome IV piRNA clusters. HSF-1::FLAG ChIP-qPCR was performed in wild-type and *hsf-1* animals under control conditions or after 30 min HS. Values are shown relative to control of each strain. *n* = 3–6 biological repeats. **d.** Heat-shock induction of the HSF-1- and PRDE-1-dependent piRNA **21ur-2348**, predicted to target *hsp-70*. Relative 21ur-2348 levels were measured in wild-type, *prde-1*, and *hsf-1(sy441)* animals before and after 30 min HS. *n* = 3–6 biological repeats. **e.** Coordinated changes in unclassified 22G-RNAs and their corresponding mRNAs during heat shock and recovery. Scatterplots show log₂ changes in 22G-RNA abundance (x-axis) versus cognate mRNA abundance (y-axis) after 30 min HS or 2 hr recovery in wild-type, *prde-1*, and *hsf-1* animals. Dashed lines indicate significance thresholds; *n* indicates the number of significant 22G-RNA/mRNA pairs. **f.** Gene Ontology enrichment of transcripts associated with the heat-shock-responsive unclassified 22G-RNA population in wild-type animals. Dot size indicates fold enrichment and color indicates adjusted *p* value; enriched terms include protein folding and chaperone-related functions. **b, c, d**: Data are mean ± SEM. Statistical analyses: two-tailed unpaired t-test (**b, c**); two-way ANOVA (**d**). ns, not significant; ✱ *p* < 0.05; ✱✱ *p* < 0.01; ✱✱✱ *p* < 0.001.

Further supporting the argument that newly induced piRNAs were responsible for generating 22G-RNAs antisense to *hsps*, almost all of the constitutively expressed piRNAs targeting *hsp-70* (*F44E5.4/.5*), the sole *hsp-70* that was induced in the HSF-1 mutants, were expressed at higher levels in the *hsf-1(sy441)* mutant (Supplementary Table 4); yet, no 22G-RNAs were produced. Thus, together these data indicate that HSF-1 activity at piRNA clusters induced new piRNA biogenesis, and was likely responsible for the production of 22G-RNAs antisense to *hsps* and potentially other heat-induced transcripts.

Because the biological functions of 22G-RNAs are largely determined by the Argonaute proteins with which they associate, we examined the Argonaute identity of the heat-induced 22G-RNAs. Unexpectedly, the HSF-1- and PRDE-1-dependent 22G-RNAs templated by *hsp* mRNAs and other heat-induced genes did not have known Argonaute associations; instead, they comprised a distinct, unclassified population^23,44^ (Fig. 2e). This ‘unclassified’ population was upregulated upon heat shock and generated from genes with molecular chaperone and protein-folding functions (Gene Ontology; adjusted p < 0.05; Fig. 2f). Other PRDE-1-dependent 22G-RNAs with known WAGO- or CSR-1-associations^23,44^ were also generated upon heat shock and were also HSF-1-dependent, but to a lesser extent; these were templated from genes enriched for ubiquitin proteasome, or cell-cycle progression and chromosome segregation functions, respectively (Supplementary Fig. 3b–e; Supplementary Table 5).

Together, these findings identify piRNA biogenesis as a previously unrecognized arm of the HSF-1-mediated heat shock response. In addition, they show that heat shock generates a distinct class of antisense 22G-RNAs, with unknown Argonaute associations, templated from newly transcribed proteostasis genes. Heat shock also generated other piRNA-dependent 22G-RNAs, also partially dependent on HSF-1, predicted to associate with WAGO- and CSR-1 Argonautes. Thus, piRNA biogenesis and the production of antisense 22G-RNAs appeared to be an intrinsic part of the *C. elegans* heat shock transcriptional program.

### The piRNA-pathway delays splicing completion

Given the established roles of piRNAs in co- and post-transcriptional silencing, we initially hypothesized that piRNA targeting of heat-induced transcripts would promote their clearance during recovery from stress; therefore, we expected *hsp* mRNAs to persist longer in *prde-1* mutants. Instead, RT-qPCR (Supplementary Fig. 4a–c) and RNA FISH (Supplementary Fig. 4d) revealed the opposite: although *hsp* mRNAs continued to accumulate for several hours after their heat-shock-triggered synthesis in both wild-type and *prde-1* mutant animals, consistent with post-transcriptional RNA processing, mRNA levels declined significantly faster in *prde-1* mutants (Supplementary Fig. 4a–c). This premature decrease was also evident by RNA FISH: *hsp-70* mRNA persisted in intestinal cells of wild-type animals even at 6 hours into recovery after heat shock exposure, but not in *prde-1* mutants (Supplementary Fig. 4d). The increased mRNA persistence in wild-type animals could not be explained by altered transcription, as Pol II occupancy levels and duration at *hsp* genes were indistinguishable between wild-type and *prde-1* animals (Supplementary Fig. 1f).

Therefore, to determine how piRNA targeting influenced the fate of newly synthesized heat-shock-induced transcripts, we pulse-labeled nascent RNA with 5-ethynyluridine (EU)^45,46^ during heat shock and quantified the conversion of labeled pre-mRNA into mature mRNA during recovery using exon-intron and exon-exon primers and absolute RT–qPCR (Fig. 3a; see Methods). Consistent with similar Pol II occupancy, wild-type and *prde-1* synthesized similar amounts of total *hsp-70* RNAs upon heat shock (Fig. 3b; Supplementary Fig. 4e). Despite that, *prde-1* mutants had significantly more EU-containing spliced mature mRNA by 2 hours of recovery from heat shock (Fig. 3c; Supplementary Fig. 4f), and less unspliced EU-labeled pre-mRNA by the end of heat shock (Fig. 3d; Supplementary Fig. 4g). For instance, for *hsp-70 (C12C8.1)*, approximately 75% of newly synthesized transcripts remained unspliced in wild-type animals after 30 min of heat shock, compared with only ∼57% in *prde-1* mutants (Fig. 3d), but *prde-1* animals had accumulated more EU-labeled spliced *hsp-70 (C12C8.1)* mRNA by 2 hr of recovery, corresponding to an approximately two fold or more higher splice-completion index 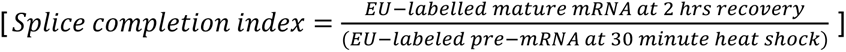 (Fig. 3e; Supplementary Table 6). This faster availability of mature *hsp-70* mRNA in *prde-1* mutants was also evident in their increased HSP-70 protein levels (Fig. 3f). For *hsp-70* (*F44E5.4/.5*), the majority of pre-mRNA was spliced by the end of the 30-minute heat shock exposure in both wild-type and *prde-1* animals; nevertheless, splice completion of the remaining pre-mRNA occurred faster in *prde-1* (Supplementary Fig. 4h; Supplementary Table 6). A similar acceleration of post-transcriptional splice completion in *prde-1* animals was also observed for *hsp-16.11* (Supplementary Fig. 4i). Thus, it appeared that one explanation for the premature depletion of mRNA in *prde-1* mutants was that accelerated splicing completion depleted pre-mRNA earlier but increased HSP-70 protein levels; in wild-type animals, an intact piRNA pathway delayed the completion of splicing and preserved nascent (EU-labeled) pre-mRNA pools, enabling continued mRNA production during recovery from stress.

**Figure 3.**
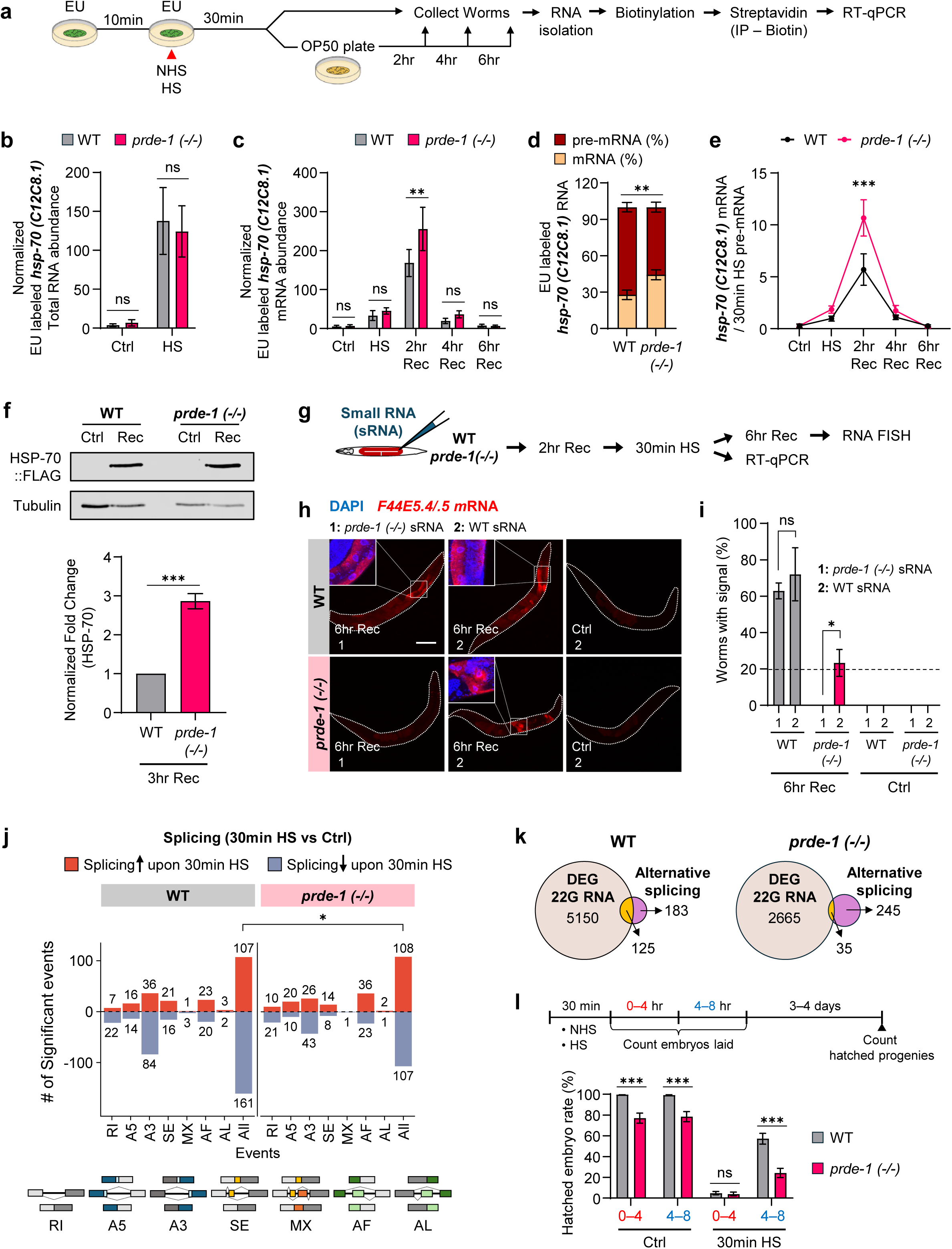
The piRNA pathway delays splice completion of heat-shock-induced transcripts. **a–e.** EU pulse-chase analysis of *hsp-70 (C12C8.1)* RNA processing. **(a)** Experimental schematic. Animals were preloaded with 5-ethynyluridine (EU) for 10 min and pulse-labeled during 30 min heat shock (HS), followed by chase on unlabeled bacteria for 2–6 hr recovery (Rec). EU-labeled RNA was biotinylated, isolated with streptavidin, and quantified by RT–qPCR, with *pmp-3* used for normalization. **(b, c)** Total EU-labeled *hsp-70* RNA (b) and mature mRNA (c) in wild-type and *prde-1* animals. **(d)** Relative proportions of EU-labeled *hsp-70* pre-mRNA and mature mRNA at the end of 30 min HS. **(e)** Ratio of EU-labeled mature mRNA to pre-mRNA during HS and recovery, showing accelerated splice completion in *prde-1*. *n* = 5–14 biological repeats. **f.** HSP-70 protein accumulation during recovery. Representative immunoblot and quantification of endogenous HSP-70 (C12C8.1)::FLAG in wild-type and *prde-1* animals 3 hr after HS. Tubulin, loading control. *n* = 3 biological repeats. **g–i.** Heat-shock-induced small RNAs restore *hsp-70* RNA persistence in *prde-1*. **(g)** Experimental schematic. < 200-nt RNA fractions isolated from heat-shocked wild-type or *prde-1* animals were microinjected before HS, and *hsp-70* RNA was assayed immediately or after 6 hr recovery. **(h)** Representative RNA FISH. *F44E5.4/.5* mRNA (red); DNA, DAPI (blue) Scale bar, 100 μm. **(i)** Percentage of injected animals retaining detectable *hsp-70* mRNA. *n* = 17–22 (6 hr Rec) and 6– 10 (control) biological repeats. **j, k.** Transcriptome-wide effects of the piRNA pathway on heat-shock-associated splicing. **(j)** Numbers of significantly increased (red) or decreased (blue) alternative splicing events following HS in wild-type and *prde-1* animals, classified as retained intron (RI), alternative 5′ or 3′ splice site (A5, A3), skipped exon (SE), mutually exclusive exons (MX), and alternative first or last exon (AF, AL). Gene models illustrate each event class. **(k)** Overlap between genes exhibiting heat-shock-responsive alternative splicing and genes generating differential 22G-RNAs in wild-type and *prde-1* animals. **l.** Recovery of reproductive capacity following heat shock. **Top**, experimental schematic; **bottom**, percentage of embryos laid during the indicated recovery intervals that subsequently hatched in wild-type and *prde-1* animals. *n* = 7 biological repeats. **b–f, i, j, l**: Data are mean ± SEM. Statistical analyses: two-way ANOVA (**b, c, e, l**); two-tailed unpaired t-test (**d, f**); Fisher’s exact test (**i**); χ² test (**j**). ns, not significant; ✱ *p* < 0.05; ✱✱ *p* < 0.01; ✱✱✱ *p* < 0.001.

To determine whether heat-induced small RNAs were sufficient to regulate the kinetics of mRNA splice completion, we microinjected the < 200-nt RNA fraction isolated from heat-shocked wild-type or *prde-1* animals into *prde-1* mutants prior to heat shock exposure and monitored their *hsp* RNA dynamics (Fig. 3g). If heat-induced 22G-RNAs delayed splice completion, wild-type—but not *prde-1—* small RNA should restore normal RNA kinetics. Remarkably, this was the case: ∼20% of *prde-1* animals injected with heat-shocked wild-type small RNAs retained *hsp-70* mRNA six hours after heat shock (RNA FISH; Fig. 3h, i), and did not how a premature decrease in the pre-mRNA pool (Supplementary Fig. 4j). In contrast, small RNAs from heat-shocked *prde-1* animals or non-heat-shocked wild-type animals had no effect (Fig. 3i). Although rescue was incomplete, likely reflecting variable RNA delivery by microinjection, these findings demonstrate that heat-induced small RNAs are sufficient to restore normal splicing kinetics.

Similar to mammalian cells, where heat shock shifts splice completion from co-transcriptional to post-transcriptional^1,9,10,47,48^ processing, the piRNA-dependent delay in splicing completion in *C. elegans* also extended beyond *hsps.* Splicing analysis using SUPPA^49^ identified 540 differentially spliced transcripts (308 genes) after heat shock in wild-type animals, compared to only 139 transcripts (280 genes) in *prde-1* (Fig. 3j). Approximately 40% (125/308) of these transcripts in wild-type animals produced antisense 22G-RNAs; only 12% (35/280) produced 22G-RNAs in *prde-1* (Fig. 3k). In addition, across all major alternative splicing (AS) events^50,51^, more wild-type transcripts retained introns and were incompletely spliced compared to *prde-1* (Fig. 3j). These delayed transcripts were enriched for developmental and neuronal functions (Supplementary Fig. 5), suggesting that the splicing delay may help re-establish developmental processes during recovery from heat shock. Accordingly, heat-shocked *prde-1* mothers exhibited a marked delay in producing viable embryos post heat-shock (Fig. 3l).

Together, these findings indicated that rather than promoting post-transcriptional silencing, the piRNA pathway prevented splicing completion of *hsp* RNAs (and other heat induced transcripts), prolonging the availability of mature mRNAs. Heat-induced, piRNA-dependent small RNA fraction alone was sufficient to restore the kinetics of splice completion and *hsp-70* mRNA persistence in *prde-1* mutants. In addition, this piRNA-dependent delay in splice completion extended beyond *hsps* to a broad set of heat-responsive transcripts.

### 22G-RNAs associate with RNA Polymerase II and MOG-7/SPP382 at the transcription complex to regulate splicing

To understand how the piRNA/22G-RNAs influenced splice completion, we tested whether 22G-RNAs interacted with nascent transcripts and the splicing machinery during transcription. For this, we adapted Photoactivatable Ribonucleoside-Enhanced Crosslinking and Immunoprecipitation (PAR-CLIP)^52^, feeding animals non-inhibitory concentrations of 4-thiouridine (4sU) (Fig. 4a; see Methods), followed by heat shock and irradiation with UV (365 nm) to crosslink newly synthesized 4sU-labeled RNAs to proteins within Pol II-transcription complexes, or used native, stringent RIP. Using these methods, Pol II, or functional, endogenously tagged spliceosomal proteins were immunoprecipitated either immediately upon 30 minutes of heat shock when *hsp* transcription was ongoing, or after 2 hours of recovery from heat shock, when Pol II was no longer engaged with the *hsp* locus (ChIP-qPCR; Supplementary Fig. 1f), but spliced *hsp* mRNAs continued to accumulate (Supplementary Fig. 4a–c).

**Figure 4.**
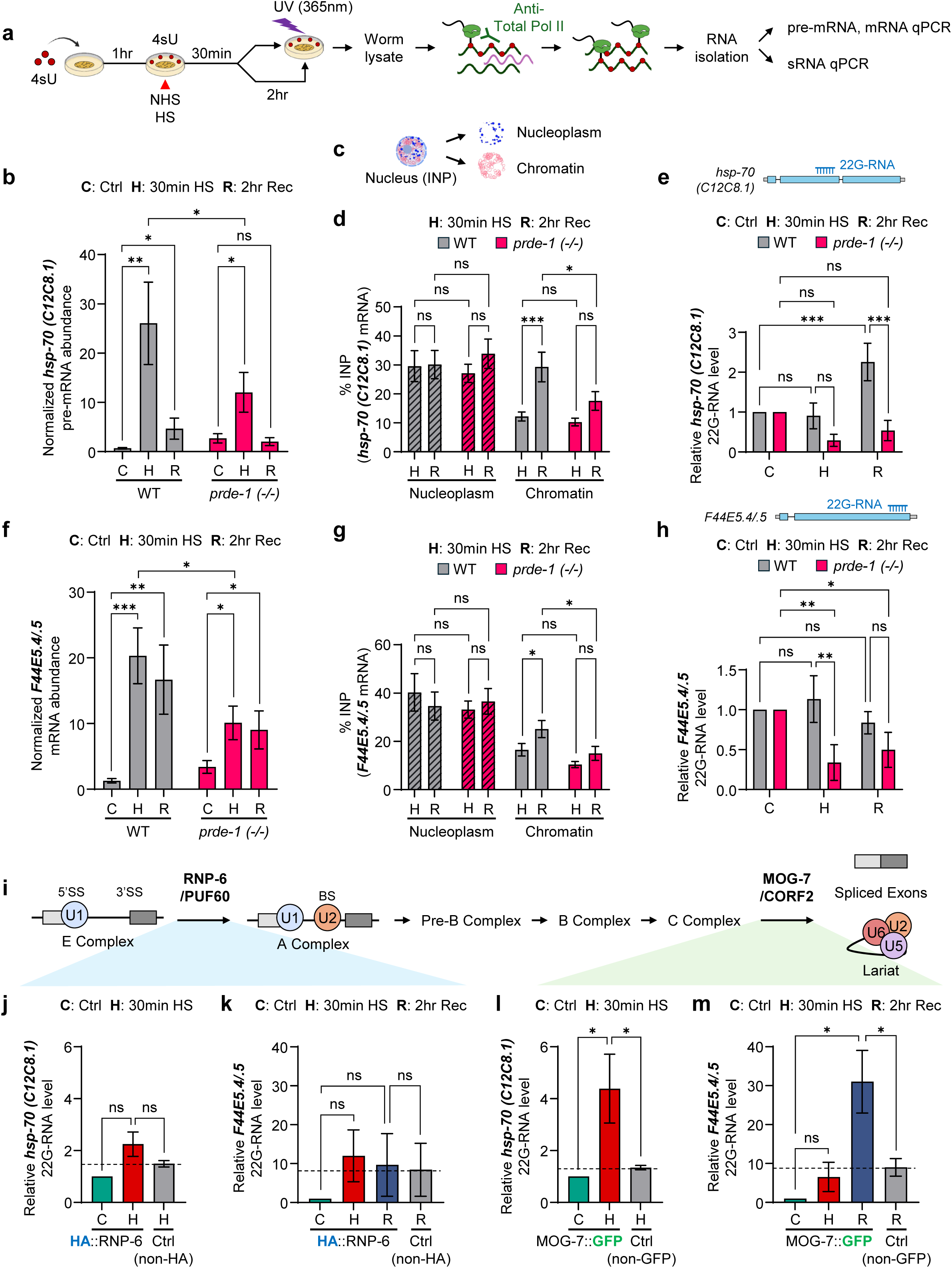
Antisense 22G-RNAs associate with RNA polymerase II and the late spliceosome factor MOG-7 during splice completion. **a–h.** Association of *hsp* RNAs and antisense 22G-RNAs with RNA polymerase II (Pol II) and chromatin. **(a)** Experimental schematic of the modified PAR-CLIP assay. Animals were fed 4-thiouridine (4sU) for 1 hr, subjected to 30 min HS, and UV-crosslinked (365 nm) immediately after HS or following 2 hr recovery. Pol II was immunoprecipitated, and associated pre-mRNAs and mRNAs were quantified by RT–qPCR and 22G-RNAs by stem-loop RT–qPCR. **(b, e)** Pol II-associated *hsp-70 (C12C8.1)* pre-mRNA (b) and antisense 22G-RNA (e) in wild-type and *prde-1* animals. **(c, d)** Experiment schematic of nuclear fractionation (c) and chromatin association of *hsp-70 (C12C8.1)* mRNA (d), expressed as percentage of nuclear input recovered in nucleoplasmic and chromatin fractions. **(f–h),** Corresponding analyses of *F44E5.4/.5*: Pol II-associated mRNA (f) nucleoplasmic and chromatin-associated mRNA (g), and Pol II-associated antisense 22G-RNA (h). Pol II-associated RNA was normalized to *pmp-3* to control for variation in immunoprecipitation. *n* = 8–13 (b, f), 9–10 (d, g), and 4–6 (e, h) biological repeats. **i–m.** Association of antisense 22G-RNAs with early and late spliceosome factors. **(i)** Simplified schematic of the spliceosome cycle showing RNP-6/PUF60 function during early spliceosome assembly and MOG-7/Ntr2/C2ORF3 during late spliceosome remodeling/discard. **(j, k)** Antisense 22G-RNAs targeting *hsp-70 (C12C8.1)* (j) and *F44E5.4/.5* (k) recovered by 4sU-treated HA::RNP-6 immunoprecipitation under control, HS, and recovery conditions; non-HA animals served as controls. **(l, m)** Antisense 22G-RNAs targeting *hsp-70 (C12C8.1)* (l) and *F44E5.4/.5* (m) recovered by native MOG-7::GFP immunoprecipitation; non-GFP animals served as controls. *N* = 4–6 (RNP-6) and 5–10 (MOG-7) biological repeats. **b, d–h, j–m**: Data are mean ± SEM. Statistical analyses: two-tailed unpaired t-test (within strain) and two-way ANOVA (between strains) (**b, f**); two-way ANOVA (within each fraction) (**d, g**); two-way ANOVA (**e, h**); Welch’s t-test (**j**–**m**). ns, not significant; ✱ *p* < 0.05; ✱✱ *p* < 0.01; ✱✱✱ *p* < 0.001.

In wild-type animals, as expected, unspliced *hsp-70* pre-mRNAs co-immunoprecipitated with Pol II at 30 minutes after heat shock, and declined during recovery (Fig. 4b). Surprisingly, newly spliced *hsp-70* mRNAs also remained associated with Pol II for at least 2 hours after heat shock (Supplementary Fig. 6a), with spliced mRNAs progressively accumulating on chromatin (Fig. 4c, d). This suggested that splice completion occurred in association with the transcription complex after transcription had ceased. Strikingly, 22G-RNAs antisense to *hsp-70* (*C12C8.1*) also co-associated with Pol II during this same recovery period (Fig. 4e), coincident with the progressive increase in spliced mRNA. Thus, 22G-RNAs were associated with Pol II at the transcription complex, and their recruitment to Pol II temporally coincides with the period of delayed splice completion.

In *prde-1* mutants, the 22G-RNAs were not recruited to the transcription complex, and the prolonged association of pre-mRNA and mRNA with Pol II was lost (Fig. 4b–e). Consistent with accelerated splice completion, less pre-mRNA remained associated with Pol II at 30 minutes heat shock (Fig. 4b). As expected, 22G-RNAs associated with Pol II were also greatly decreased (Fig. 4e), and newly spliced *hsp-70* mRNA failed to accumulate on chromatin (Fig. 4c, d), or in association with Pol II above basal levels (Supplementary Fig. 6a). These findings, together with observation that similar amounts of RNAs were synthesized upon heat shock in both *prde-1* and wild-type animals (Fig. 3b), indicated that, in the absence of piRNAs, transcripts complete splicing and disengage from the transcription complex more rapidly.

A similar overall pattern was observed with *hsp-70 (F44E5.4/.5)*, although the majority of this transcript was already spliced by the end of 30-minute heat shock, in both wild-type and *prde-1* (Supplementary Fig. 4g). Here too, by 30 minutes more antisense 22G-RNAs associated with Pol II in wild-type animals than *prde-1* mutants (Fig. 4h), as did more mature mRNA (Fig. 4f), and mRNA continued to accumulate on chromatin during the 2-hour recovery (Fig. 4g) after transcription had ceased, indicative of post-transcriptional splicing. Thus, although 22G-RNA association with Pol II differed somewhat between the two *hsp-70* genes, consistent with their distinct splicing kinetics (Supplementary Fig. 4a–c; Fig 3d; Supplementary Fig. 4g), both transcripts exhibited the same fundamental relationship: piRNA-generated 22G-RNAs associated with Pol II during the course of pre-mRNA splicing into mature mRNA.

Association with spliceosomal proteins further supported the model that 22G-RNAs influenced splicing at the transcription complex. To assess 22G-RNA interactions, we chose two spliceosome proteins that function at distinct stages of the splicing cycle: functional, endogenous HA-tagged RNP-6^53,54^, the *C. elegans* ortholog of mammalian PUF60, which promotes early spliceosome assembly through recognition of polypyrimidine tracts and U2 snRNP recruitment, and functional GFP-tagged MOG-7^55^, the ortholog of the yeast Nineteen complex-related protein Ntr1/Ntr2/SPP382, an interacting partner of the PRP43 helicase which facilitates intron lariat release and assists in splicing quality control, later in the splicing cycle^56–59^ (Fig. 4i). Associated 22G-RNAs were quantified by stem-loop RT-qPCR, with co-immunoprecipitated pre-mRNAs serving as a positive control for RNA–protein crosslinking.

Although *hsp* pre-mRNAs co-immunoprecipitated with RNP-6 (Supplementary Fig. 6b, c), antisense 22G-RNAs were not detectably enriched above background levels (Fig. 4i–k), suggesting that these small RNAs are not associated with early spliceosome assembly. In contrast, antisense 22G-RNAs co-immunoprecipitated with the late spliceosome factor MOG-7 together with *hsp* pre-mRNAs, consistent with a role in late spliceosomal processes and splicing quality control (Fig. 4i, l, m; Supplementary Fig. 6d, e). For reasons we do not yet understand but that may be related to assay resolution, splicing mechanisms regulated by MOG-7, and the splicing kinetics and intron numbers of the *hsp* genes, the timing of 22G-RNA association with MOG-7 differed between the two *hsp-70s*. *hsp-70* (*C12C8.1*)-targeting-22G-RNAs associated with MOG-7 during the 30-minute heat shock (Fig. 4l), whereas 22G-RNAs antisense to *hsp-70* (*F44E5*.4/.5) were associated with MOG-7 primarily during the 2-hour recovery period (Fig. 4m). Wild-type animals lacking tagged proteins served as negative controls, confirming the specificity of these interactions.

Together, these findings place piRNA-generated 22G-RNAs at the transcription complex during the interval of delayed splice completion and link them to late spliceosome remodeling and quality control events. In wild-type animals, 22G-RNAs co-associated with Pol II as nascent transcripts continued to undergo splicing and accumulate in chromatin after transcription had ceased; in *prde-1* mutants, these associations were lost, and transcripts completed splicing and disengaged from Pol II more rapidly. The association of 22G-RNAs with the late spliceosome quality-control factor MOG-7 further supports a role for these endo-siRNAs in splicing regulation. Thus, these data support a model whereby piRNA-generated 22G-RNAs act locally at late stages of the splicing cycle to restrain splice completion and prolong transcript association with the transcription machinery.

### The piRNA pathway preserves transcript fidelity

Given the association of 22G-RNAs with spliceosome quality control, we considered the possibility that delayed splicing, *via* the piRNA system, may also serve a broader function in guaranteeing splicing fidelity during the rapid transcriptional changes that accompany heat shock. To test whether this was the case, we sequenced mRNAs from heat shock-exposed wild-type and *prde-1* mutants after incorporating unique molecular identifiers (UMIs)^60–64^ into individual mRNAs during first-strand cDNA synthesis so as to distinguish genuine transcript variants from PCR or sequencing artifacts and analyzed the RNA for transcript variants or errors (Supplementary Fig. 7a; see Methods).

When analyzed using two methods—genome-wide mismatch frequencies and stringent variant calling (GATK HaplotypeCaller^65,66^; see Methods)— mRNA from *prde-1* mutants consistently contained higher transcript variants or ‘errors’ than wild type (Fig. 5a, b; Supplementary Tables 7 and 8). The majority were base substitutions, with G>A transitions predominating, consistent with increased misincorporation of nucleotides by Pol II^67^ (Supplementary Fig. 7b). In addition, the *prde-1* variant mRNAs were not uniformly distributed across the genome; instead, they were disproportionately derived from smaller genes, which were usually more highly expressed (Fig. 5b). Thus, it appeared that the piRNA-dependent delay in splice completion might promote transcript fidelity during stress, and *prde-1* mutants accumulated a modest but significant increase in mRNA variants.

**Figure 5.**
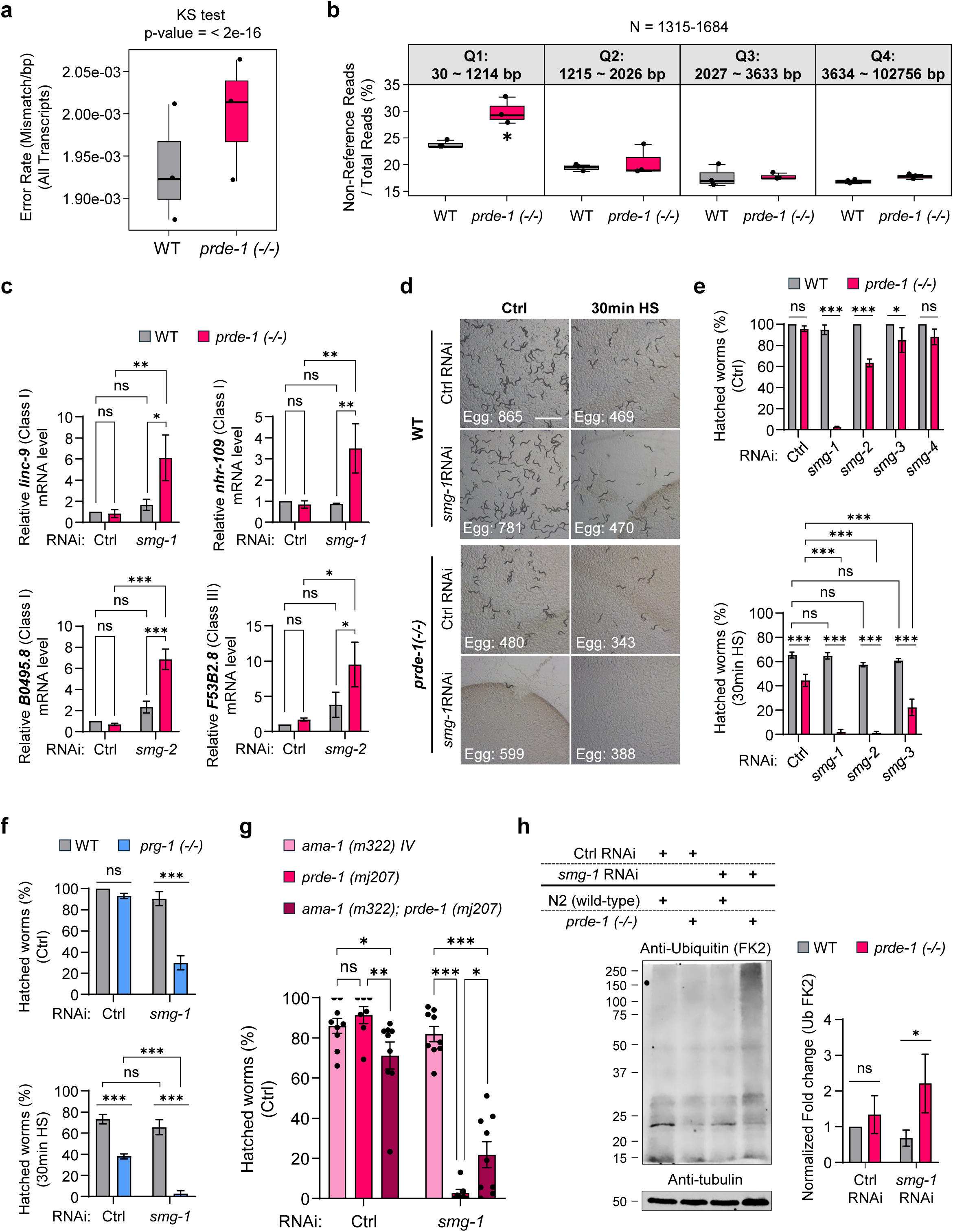
piRNA protects transcript fidelity and complements nonsense-mediated decay (NMD) to maintain protein quality control. **a, b.** Transcript variants in wild-type and *prde-1* animals. **(a)** Transcriptome-wide mismatch frequency determined from UMI-tagged RNA sequencing. Approximately 3.71 × 10^7 individual mRNAs were analyzed per biological replicate and genotype. **(b)** Percentage of non-reference reads stratified by gene length quartile (Q1, shortest; Q4, longest) in wild-type and *prde-1* animals. Numbers above indicate the gene-length range and number of genes within each quartile. **c.** Accumulation of validated NMD substrates following NMD inhibition. RT–qPCR of Class I (*linc-9, nhr-109,* and *B0495.8*) and Class III (*F53B2.8*) NMD targets in wild-type and *prde-1* animals treated with control, *smg-1*, or *smg-2* RNAi. RNA levels were normalized to *eft-3* and expressed relative to control RNAi-treated wild type. *n* = 4–6 biological repeats. **d–f.** Dependence of piRNA-pathway mutants on NMD for embryonic viability. **(d)** Representative images of progeny from control or heat-shocked wild-type and *prde-1* animals treated with control or *smg-1* RNAi; numbers indicate eggs laid in the representative experiment. Scale bar, 2 mm. **(e)** Percentage of hatched progeny from wild-type and *prde-1* animals following RNAi against the indicated NMD components under control (top) or heat-shock (bottom) conditions. **(f)** Embryonic viability of wild-type and *prg-1* animals following control or *smg-1* RNAi, under control (top) or heat-shock (bottom) conditions. *n* = 3–5 (e) and 4–8 (f) biological repeats. **g,** Slowing RNA polymerase II elongation partially suppresses the NMD dependence of *prde-1*. Embryonic viability of *ama-1(m322)*, *prde-1(mj207)*, and *ama-1(m322); prde-1(mj207)* animals following control or *smg-1* RNAi. Points represent independent biological experiments (*n* = 7–9). **h,** Loss of NMD increases the burden of polyubiquitinated proteins in *prde-1*. Representative FK2 immunoblot (left) and quantification (right) of polyubiquitinated proteins in wild-type and *prde-1* animals following control or *smg-1* RNAi. Tubulin, loading control. *n* = 5 biological repeats. **a, b**: Data are median ± SEM. Statistical analyses: Kolmogorov–Smirnov test (**a**); two-tailed unpaired t-test (**b**). ✱ *p* < 0.05. **c, e–f**: Data are mean ± SEM. Statistical analyses: two-way ANOVA. ns, not significant; ✱ *p* < 0.05; ✱✱ *p* < 0.01; ✱✱✱ *p* < 0.001.

The increase in aberrant transcripts in *prde-1* animals led us to examine whether *prde-1* animals had upregulated the nonsense-mediated mRNA decay (NMD) pathway, a system that surveils and eliminates unproductively spliced transcripts^35,68–72^. This would be reflected by a greater increase in mRNAs normally degraded by NMD in *prde-1* animals than in wild-type following NMD inhibition. Indeed, this appeared to be the case: four previously validated *C. elegans* NMD targets^72^ were upregulated to a greater extent in *prde-1* than in wild-type animals when *smg-1,* which encodes the NMD kinase, or *smg-2,* the *C. elegans* homolog of UPF1^35,73^, were knocked down using RNA interference (Fig. 5c).

Remarkably, *prde-1* animals not only exhibited elevated NMD activity, but they were also profoundly reliant on NMD for embryonic viability. Under both control conditions and following heat shock, *smg-1* knockdown caused complete embryonic lethality in *prde-1* animals (Fig. 5d, e). Likewise, knockdown of *smg-2* (UPF1 homolog) or *smg-3* (UPF2 homolog)^73^ in *prde-1* mutants also caused a modest but significant loss of embryonic viability under control conditions, and severe loss of embryonic viability upon heat-shock (Fig. 5e). In contrast, wild-type animals did not display this synthetic lethal phenotype, and embryos remained fully viable following *smg-1*, *smg-2*, or *smg-3* RNAi under both control and heat-shock conditions (Fig. 5d, e), consistent with previous reports that NMD is dispensable for normal *C. elegans* development^35,71–74^. A similar phenotype was observed in *prg-1* mutants, confirming the NMD dependence of piRNA pathway deficiency (Fig. 5f). Remarkably, crossing *prde-1* with a slow Pol II elongation mutant, *ama-1(r739h)*^75,76^, that decreases the baseline speed of transcription and is predicted to decrease splicing kinetics, partially rescued the synthetic lethality caused by *smg-1* downregulation in *prde-*1, although the extent of rescue varied between populations (Fig. 5g).

Consistent with the loss of the piRNA pathway increasing transcript errors, in the absence of *smg-1*, *prde-1* animals accumulated significantly higher amounts of polyubiquitinated proteins even under control conditions in the absence of heat shock exposure (Fig. 5h). Together, these data indicated that the loss of the piRNA system was generating transcripts with transcriptional or splicing errors that required elimination by NMD (AS-NMD^77–80^) in order to preserve embryonic viability and prevent the accumulation of misfolded proteins.

### The piRNA pathway preserves splice-site selection and suppresses the formation of NMD-sensitive transcripts

To identify the splicing defects that arise from the loss of piRNAs, we performed long-read RNA sequencing on wild-type and *prde-1* mutants after inhibiting nonsense-mediated decay (NMD) using *smg-2* RNAi (Supplementary Fig. 8a, b; Supplementary Table 9). This preserved aberrantly spliced transcripts that would otherwise have been rapidly degraded by NMD^77–80^. As in previous experiments, day-one adults, which are viable upon NMD inhibition but accumulate transcript variants and protein aggregates upon loss of the piRNA pathway, were examined.

Strikingly, the three highly induced *hsp*s, which underwent faster splice completion in *prde-1* mutants, displayed strong use of the expected junctions in wild-type animals, but exhibited greater heterogeneity in junction choice and increased presence of multiple splice junctions in *prde-1* mutants, consistent with less constrained splice-site selection (Fig. 6a; Supplementary Fig. 8c, d). These events generally became most prominent in *prde-1* upon *smg-2* depletion, also supporting the model that the loss of piRNA-dependent 22G-RNA production increased NMD dependence due to the synthesis of aberrantly spliced transcripts.

**Figure 6.**
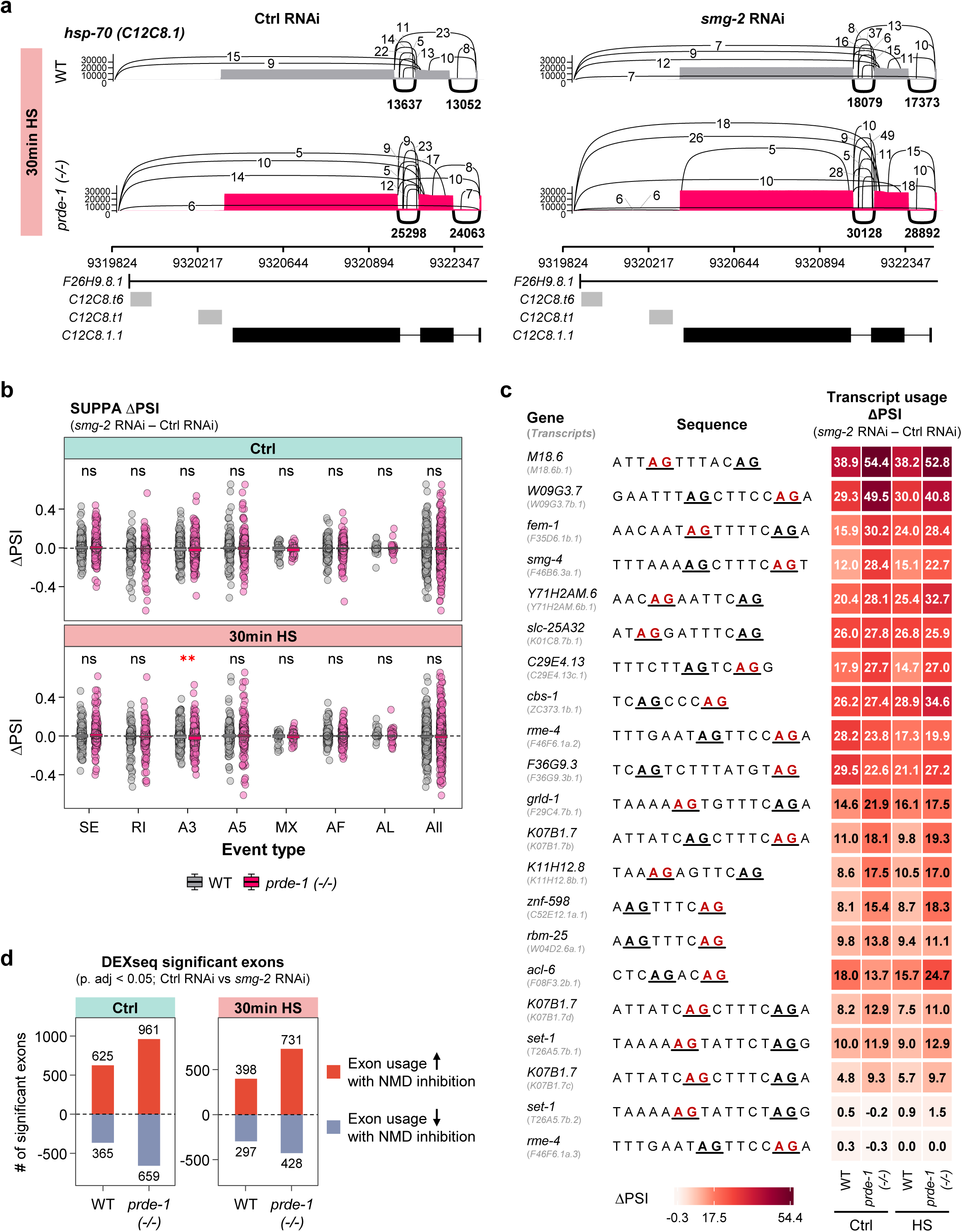
The piRNA pathway ensures splice-site and exon usage to limit the production of NMD-sensitive isoforms. **a.** Sashimi plots of *hsp-70 (C12C8.1)* after 30 min heat shock in wild-type and *prde-1* animals treated with control or *smg-2* RNAi. Curves and numbers indicate splice junctions and junction-supporting reads, respectively. Annotated transcript models below include *F26H9.8.1 (minus strand)*, *C12C8.t6*, *C12C8.t1*, and *C12C8.1.1.* **b–d.** Alternative splicing events revealed by NMD inhibition (AS-NMD) in wild-type and *prde-1*. **(b)** SUPPA2 analysis showing ΔPSI (*smg-2* RNAi − control RNAi) for skipped exon (SE), retained intron (RI), alternative 3′ or 5′ splice site (A3, A5), mutually exclusive exon (MX), alternative first exon (AF), and alternative last exon (AL) events under control and heat-shock conditions. Each point represents an individual event; wild-type (gray), *prde-1* (magenta). Adjusted Wilcoxon rank-sum P values compare genotypes within each event class. **(c)** Alternative 3′ splice-site usage among NMD-sensitive transcripts from genes exhibiting A3 events in both wild-type and *prde-1*. Competing canonical AG acceptors are shown in the sequence, with the NMD-associated acceptor in red. Heatmap shows the change in transcript usage following NMD inhibition (*smg-2* RNAi − control RNAi) in each genotype and condition. **(d)** DEXSeq analysis of differential exon usage following NMD inhibition. Bars indicate numbers of exons with significantly increased (red) or decreased (blue) usage (p. adj < 0.05) in wild-type and *prde-1* animals under control and heat-shock conditions.

To determine whether specific classes of alternative splicing events were preferentially affected by loss of the piRNA pathway, we quantified differential splicing events using SUPPA (adjusted p < 0.05). Consistent with our short-read RNA-seq analysis (Fig. 3j), long-read sequencing also showed that, with NMD intact, wild-type animals exhibited substantially more significant alternative splicing events than *prde-1* mutants under both basal and heat-shock conditions (129 versus 51 events in control; 119 versus 55 following heat shock; Supplementary Fig. 9a). However, when NMD was downregulated by *smg-2* RNAi, significantly more alternative splicing events became evident in *prde-1* mutants than in wild-type animals (Supplementary Fig. 9b; Supplementary Table 10), indicating that *prde-1* animals normally produced more unproductively spliced transcripts that were then eliminated by NMD (AS-NMD). For instance, SUPPA analyses identified 179 vs.129 total significant AS-NMD-sensitive events in *prde-1* versus wild-type upon heat shock (Supplementary Fig. 9b). Moreover, although NMD inhibition increased the abundance of all major classes of alternative splicing events in both wild-type and *prde-1* animals, indicating that, as in mammals, *C. elegans* normally generates several AS-NMD isoforms, alternative 3′ splice-site (A3) events showed the clearest difference between wild type and *prde-1* (Fig. 6b).

To determine whether this difference in A3 events between *prde-1* and wild-type was due to altered splice-site choice or splice-site recognition, we examined the consensus 3’ acceptor sequences^81^ (and 5’ donor sequences) used by wild-type and *prde-1* animals. These consensus sequences were nearly identical (Supplementary Fig. 9c), suggesting that the loss of the piRNA pathway did not affect 3’-splice-site recognition. Rather, it altered splice-site selection between competing canonical AG dinucleotide acceptor sequences: when A3 sites in 19 events shared by both wild-type and *prde-1* animals were examined, in *prde-1* animals, splice-site usage was shifted in favor of those that generated isoforms normally targets of NMD (Fig. 6c; ΔPSI (*smg-2* RNAi – Control RNAi) values are higher in *prde-1* under control and heat shock conditions; also see Supplementary Fig. 9d). Thus, in wild-type animals, the piRNA pathway biased A3 choice to suppress the generation of NMD-sensitive splice variants; in *prde-1* mutants, this stringency of splice site selection was lost.

A second major class of alternatively spliced mRNAs that were stabilized by NMD inhibition were those exhibiting differential abundance of individual exons relative to other exons within the same transcript (DEXSeq)^82^. This suggested that differential exon usage was also a signature of AS-NMD in *C. elegans,* and these transcripts too were significantly and markedly more abundant in *prde-1* than in wild-type animals (Fig. 6d; Supplementary Table 11). Notably, among these were mRNAs from genes encoding spliceosomal and RNA-processing factors, including *hel-1* and *rsp-1*^83^ (Supplementary Fig. 10a, b), raising the possibility that altered splicing of core splicing regulators further amplifies transcriptome-wide effects on splicing caused by the loss of piRNA pathway. Consistent with this, IsoQuant reconstruction^84^ of full-length transcripts revealed a corresponding increase in altered transcript architectures in *prde-1* animals compared to wild-type following NMD inhibition (Supplementary Fig. 10c).

Together, these data indicate that the piRNA pathway preserves the fidelity of splicing. Particularly, evident splicing defects upon the loss of the piRNA pathway were altered 3’ splice-site choice and exon usage: in *prde-1* animals, the spliceosome more frequently committed to an alternative 3’ splice site or included or omitted exons that favor the production of transcript isoforms that are normally eliminated by NMD. Combined with the widespread changes in transcript architecture that occur upon the loss of the piRNA pathway, and our biochemical evidence that piRNA-generated 22G-RNAs associate with the transcription complex and the late spliceosomal quality-control factor MOG-7, while delaying splice completion, these findings suggest that the piRNA-dependent delay in splice completion might preserve accurate splice site selection, thereby maintaining transcript isoform fidelity.

## Discussion

The splice site sequences in higher eukaryotic introns are highly degenerate and alone do not contain sufficient information to accurately determine bona fide splice-site usage, or which isoform needs to be produced^1,85–88^. Thus, how cells regulate splice-site selection and splicing fidelity remains poorly understood. Here, we identify an unexpected role for the piRNA pathway in this process. By leveraging the heat shock response to follow the role of piRNA pathway in gene expression regulation, we show that during heat shock, piRNA-triggered sequence-specific antisense 22G-RNAs generated from *hsp* transcripts interact with Pol II and the late spliceosome protein MOG-7/Ntr1/C2ORF3 at the transcription complex to delay splice completion and regulate splice site usage. Surprisingly, the heat shock transcription factor HSF-1, which not only triggers the expression of *hsp* genes and other protective mRNAs, also induces piRNAs upon heat shock, and at least one of them is capable of targeting the newly transcribed *hsp* mRNAs to generate ‘splice-assisting’ 22G-RNAs. Together, these findings implicate the piRNA pathway in splicing regulation and suggest that HSF-1 coordinates stress-induced transcription with the splicing fidelity of the transcripts it produces. This mechanism is reminiscent of HSF1 activity in mammalian cells, where HSF1 induced Satellite III (SatIII) repeats generate noncoding RNAs that recruit RNA-processing and splicing factors to nuclear stress bodies, coupling transcriptional activation to noncoding RNA-mediated control of splicing^89,90^.

How piRNAs influence widespread splice-site selection remains unclear. Our data establish the germline origin of piRNA targeting but do not determine how the resulting 22G-RNAs access transcripts more broadly expressed, likely across the animal. Whether 22G-RNAs themselves move between tissues, engage other systemic mechanisms triggered by piRNAs and PIWI/PRG-1, or interact with other Argonaute proteins remains unknown^91–96^. Also, while we leveraged the unique advantages of heat shock and the inducible expression of *hsp* genes to gain mechanistic insight into piRNA-dependent splicing regulation, long read sequencing revealed that the loss of the piRNA pathway regulates splicing beyond *hsps*, altering exon and alternative 3’ splice-site usage of several genes to bias transcript expression to favor isoforms normally eliminated by NMD. This implies that under normal circumstances, in wild-type animals, the piRNA pathway suppresses splicing at these sites, minimizing the production of unproductively spliced transcripts that require clearance by NMD. The association of piRNA-generated 22G-RNAs with MOG-7/NTR spliceosome-remodeling machinery, and the antisense nature of these small RNAs, provide an attractive plausible mechanism for conferring sequence specificity on splice-site selection, and raise the possibility that sequence-guided regulation acts during late stages of splicing quality control. However, the detailed mechanisms of all of these steps remain to be worked out.

A striking consequence of the loss of the piRNA pathway in *C. elegans* is a profound sensitivity to inhibition of NMD. NMD impairment results in the accumulation of misfolded proteins and synthetic embryonic lethality, even in the absence of heat shock, again indicating that piRNA-deficient animals likely continuously generate transcripts that require RNA surveillance and elimination to ensure development and physiology. Remarkably, and consistent with the kinetic model of splicing^97,98^, slowing Pol II elongation partially restored embryonic viability to piRNA-deficient animals upon NMD knockdown. Together, these findings provide further support for the role of the piRNA pathway in RNA quality. More broadly, these findings raise the intriguing possibility that, analogous to therapeutic splice-switching antisense oligonucleotides, *C. elegans* has evolved an endogenous small RNA-guided mechanism to ensure accurate splicing and splice-site selection across its highly compact, nested, gene-dense genome.

What might be the rationale for this unexpected role of piRNAs in splicing? Ancestrally piRNAs target transposable elements (TEs) in the germline of most animals, suppressing their mobilization to preserve genome integrity. However, TEs are also considered major drivers of intron evolution, having repeatedly inserted themselves into genes to introduce cryptic splice sites, branch points, polypyrimidine tracts, and splicing regulatory motifs^33,99–102^. In humans, a significant proportion of recently acquired splice signals are located within intronic transposable elements^103,104^. Likewise, splicing junctions between exons and TEs are thought to result in TE exonization—mature chimeric mRNAs that incorporate TE-derived sequences— and are frequently eliminated by NMD^103,104^. Thus, an attractive possibility is that the piRNA pathways’ role in splicing regulation evolved from its ability to detect these TE insertion-derived sequence “scars”, and route them to NMD to preserve splicing fidelity. Such a model could provide a conceptual link between the canonical role of piRNAs in genome defense and their unexpected function in splicing quality control uncovered here. Indeed, alternative splicing has been described as a form of ‘evolutionary tinkering’^100,101^ that evolved as an inevitable consequence of splicing errors and transposition events, later selected to result in the structural and functional diversification of proteins.

Very few previous studies have linked the piRNA pathway to splicing regulation^105,106^. Yet, these studies suggest that the piRNA-splicing-regulation may be conserved. Thus, in *Drosophila*^105^ nuclear Piwi complexes inhibit splicing of the *gypsy* retrotransposon through chromatin-based, co-transcriptional silencing, and in locusts^106^ piRNAs promote splicing of specific pre-mRNAs during oocyte maturation. Particularly relevant to our findings, Barberán-Soler and colleagues^107^ showed that the PRG-1/piRNA pathway regulates alternative 3′ splice-site selection of the *C. elegans* TOR ortholog, *let-363*, in the male germline through recognition of an intronic Helitron transposon. In addition, introns protect transcripts from default Argonaute-mediated silencing in *C. elegans*, and the nuclear Argonaute HRDE-1 interacts with the conserved RNA helicase Aquarius/EMB-4, further implicating the piRNA pathway in RNA processing^108–110^. Finally, recent studies in mammals show that piRNA biogenesis is coupled to the pioneer round of translation and that piRNAs are enriched for stop codon-containing sequences^111^, linking the pathway to early RNA quality-control mechanisms.

In summary, our data show that piRNAs target mRNAs to generate 22G-RNAs in the perinuclear nuage, where decisions regarding mRNA localization, NMD, and translation are also regulated; subsequently, the 22G-RNAs control splicing completion and guide accurate splice-site selection of target genes in the nucleus in association with the transcription complex, Pol II, and late splicing and splicing quality control complex MOG-7, after transcription had ceased. Such a bidirectional mechanism positions the piRNA pathway as a liaison between the major steps of gene regulation—transcription, splice completion, RNA quality control, and protein biogenesis.

## Acknowledgements

We thank Ms. Selvarasu for microinjection and RT-qPCR experiments, the V.P lab members, and Drs. Eugene Kandel, Katherina Gurova, Andrei Gudkov, Josep Comeron, Anna Malkova, and Ana Llopart for comments, Rochester Genomics Research Center for small RNA sequencing and advice, Drs. Paul Quinn and Prashant Singh for mRNA sequencing, UMI libraries, and advice; James Coughlin and Bogdan Sieriebriennikov at Weill Cornell Medicine for advice and help with PacBio HiFi sequencing; and the Caenorhabditis Genetics Center (CGC) (funded by the NIH Infrastructure Programs P40 OD010440) for *C. elegans* strains. We thank Dr. Adam Antebi and Dr. Roger Pocock for the tagged RNP-6 and MOG-7 strains. This work was supported by NIH R01 MH126282 (V.P.), and support for the sequencing core from the National Cancer Institute (NCI) grant P30CA016056 to the Roswell Park Comprehensive Cancer Center.

## *C. elegans* STRAINS AND GROWTH CONDITIONS

*C. elegans* used in all experiments are listed in Method Table 1. Strains were obtained from the *Caenorhabditis Genetics Center* (CGC, Twin Cities, MN), generated in the Prahlad laboratory, kind gifts of Dr. Adam Antebi (Max Planck Institute for Biology of Aging), or Dr. Richard Pocock (Monash Biomedicine Discovery Institute), or generated by SunyBiotech (Suzhou, Jiangsu, China). All crossed worms were verified by genotyping. The primers used in genotyping are listed in Method Table 2.

**Table 1.** Strain information.

| Strain name | Genotype |  | Source |
| --- | --- | --- | --- |
| N2 | Bristol N2; wild-type |  | CGC |
| SX2499 | *prde-1 (mj207) V* |  | CGC |
| PS3551 | *hsf-1 (sy441) I* |  | CGC |
| FLAG::HSF-1 | *3X FLAG::hsf-1 I* | CRISPR /Cas-9 | Prahlad Lab |
| FLAG::HSF-1<br>(*sy441*) | *3X FLAG::hsf-1 (sy441) I* | CRISPR /Cas-9 | Prahlad Lab |
| FLAG::HSF-1;<br>PHX4493 | *3X FLAG::hsf-1 I; syb4493 [HA::prde-1] V* | CRISPR/Cas-9<br>Cross | Prahlad Lab /<br>SunnyBiotech |
| SX922 | *prg-1 (n4357) I* |  | CGC |
| WM274 | *prg-1 (tm872) I; neSi14 [FLAG::prg-1 + Cbr-unc-119(+)] II; unc-119 (ed3) III* |  | CGC |
| WM274;<br>SX2499 | *prg-1 (tm872) I; neSi14 II; unc-119 (ed3) III; prde-1(mj207) V* | Cross | Prahlad Lab |
| PHX2141 | *syb2141 [C12C8.1::FLAG] I* | CRISPR /Cas-9 | Prahlad Lab /<br>SunnyBiotech |
| PHX2141;<br>SX2499 | *syb2141 I; prde-1(mj207) V* | Cross | Prahlad Lab |
| DR786 | *ama-1 (m322) IV* |  | CGC |
| DR786;<br>SX2499 | *ama-1 (m322) IV; prde-1 (mj207) V* | Cross | Prahlad Lab |
| AA4624 | *dh1188 [HA::rnp-6] I* |  | Antebi Lab |
| RJP4513 | *rp127[mog-7::Degron::GFP] I; unc-119(ed3) III; ieSi38[sun-1p::TIR1::mRuby::sun-1 3'UTR + Cbr-unc-119(+)],ItIs37[(pAA64) pie-1p::mCherry::his-58 + unc-119(+)] IV; ItIs38[sun-1p::TIR1::mRuby::sun-1 3'UTR + Cbr-unc-119(+)] IV* |  | Pocock Lab |

**Table 2.**
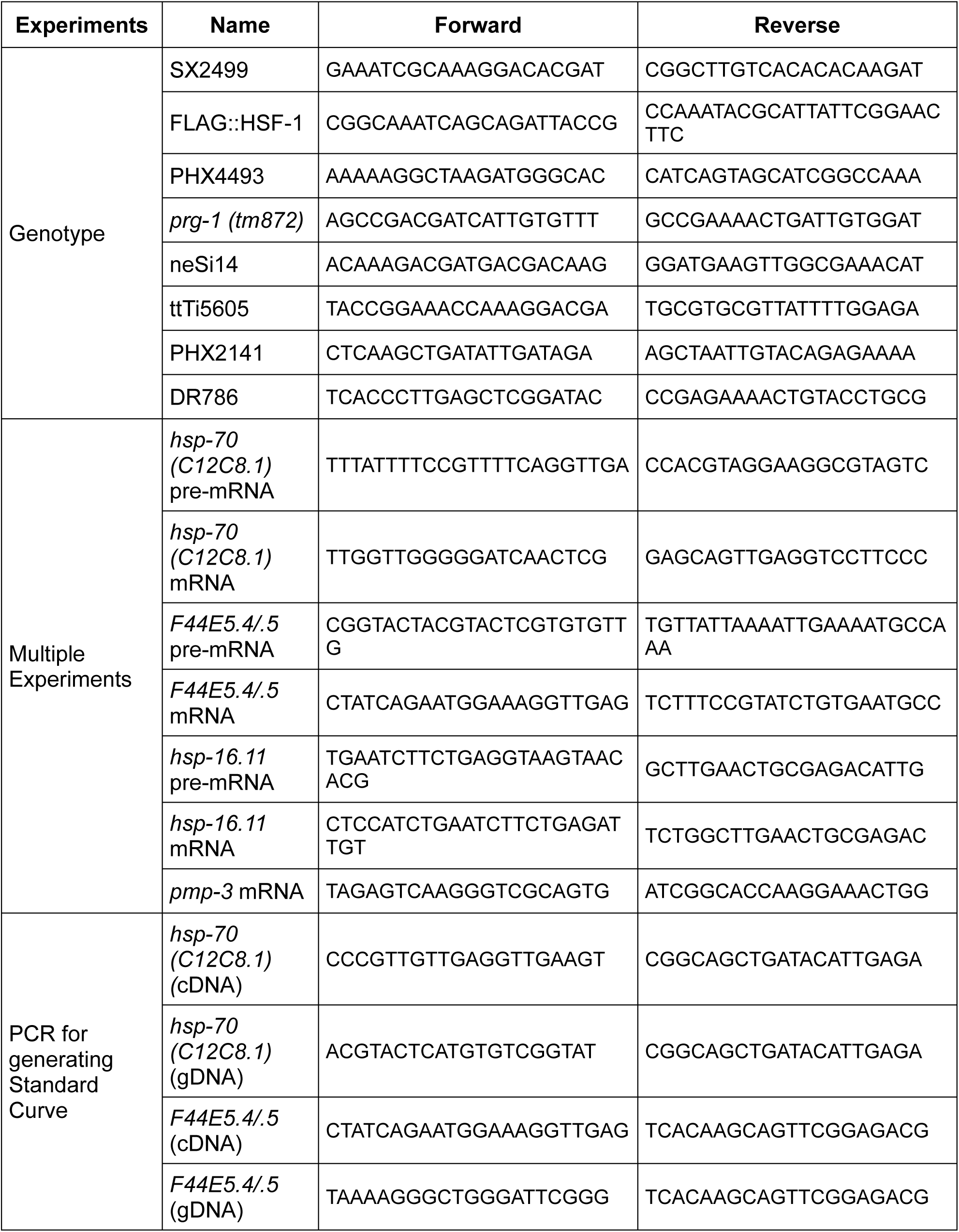
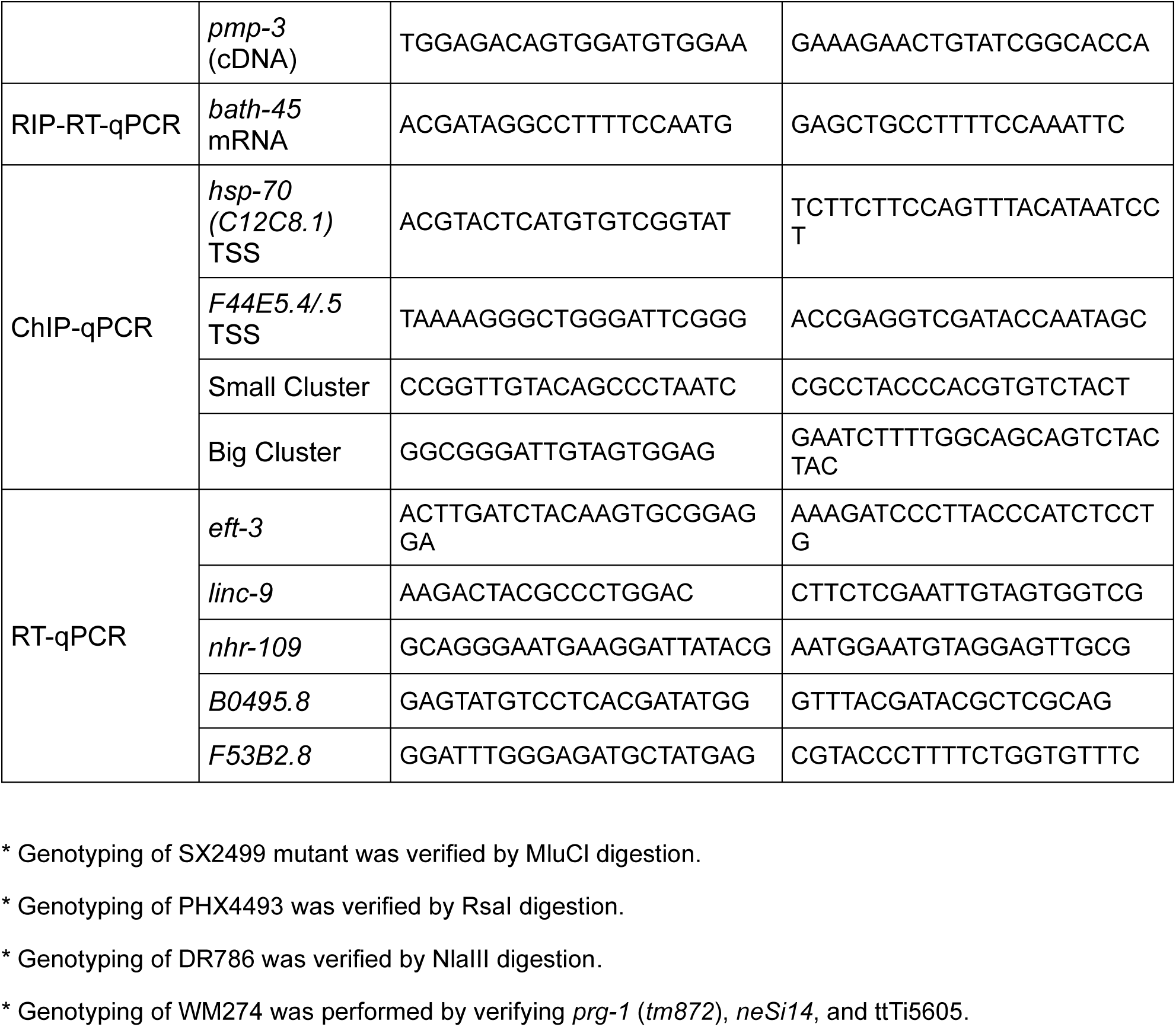
Primer information.

All strains were maintained at 20°C on nematode growth media (NGM) plates. NGM plates were prepared by pouring 8.9 ml of autoclaved liquid NGM per 60 mm plate. NGM was allowed to solidify at room temperature (RT) for 2 days, seeded with 300 μl OP50 E. coli (O.D = 1.8–2.0), and dried at RT for no more than 2 days. Population densities were matched based on growth rate of the strains. To maintain healthy, continuously growing populations, 10–15 animals at larval stage 4 (L4) were transferred to a new plate every 4 days. For experiments, day-one adult animals were used. These were generated either by picking L4-stage worms and harvesting them 24 hr later, or by bleach-hatching and collecting day-one adult worms 82–84 hr later, depending on experimental requirements.

## HEAT SHOCK

Worms were exposed to acute heat stress as previously described in Das et al., eLife 2020^1^. Briefly, NGM plates containing day-one adults were sealed with parafilm and immersed in a water bath pre-warmed to 34°C for different durations as mentioned. For recovery experiments, NGM plates were incubated in a 20°C incubator immediately after heat shock and harvested after specific times mentioned in the experiments.

## RNA SEQUENCING

### ILLUMINA SEQUENCING

#### RNA Library Preparation

For RNA-sequencing, three independent biological replicates of each strain and each treatment were harvested and processed as described. For each sample/treatment, approximately 200 L4-stage worms were transferred to OP50 plates. After 26–27 hr, worms from two plates per condition [non-heat shock (NHS), 5min HS, 30min HS, and 30min HS + 2hr Rec] were collected in nuclease-free water. Total RNA was extracted using the Mirvana miRNA Isolation Kit (Invitrogen; Cat #AM1560). Genomic DNA in the sample was removed using Turbo DNA-free Kit (Invitrogen; Cat #AM1907). RNA was separated by size into mRNA (> 200 nt) and small RNA (< 200 nt). Small RNA samples were treated with 10 U RppH (NEB; Cat #M0356S) at 37°C for 30 min, and then purified using the Zymo RNA Clean & Concentrator kit (Zymo Research; Cat #R1015). RNA concentrations were measured with a Qubit 4 fluorometer (Thermo Scientific). mRNA libraries were prepared using the KAPA mRNA Hyperprep kit (Roche; Cat #08098123702); small RNA libraries were prepared using the Illumina TruSeq Small RNA Library Preparation kit (Illumina; Cat #RS-200), following manufacturer’s protocol.

#### Small RNA analysis

Small RNA data was analyzed using the tinyRNA^2^ (v1.50) pipeline with its default configuration. The process included adapter trimming and quality filtering with fastp^3^, followed by genome alignment to the *C. elegans* genome (release WS279) using Bowtie^4^. An additional setting, counter decollapse: True, was specified to generate de-collapsed SAM files.

The module tiny-count was used to count the different types of small RNA according to the selection rules specified in the following table.

| Select for... | with value... | Classify<br>as... | Hiera<br>rchy | Overlap | Strand | 5'End<br>Nucleotide | Length |
| --- | --- | --- | --- | --- | --- | --- | --- |
| Class | risiRNA | rRNA | 1 | Partial | both | all | all |
| Class | miRNA | miRNA | 2 | 5'Anchored, 0,<br>4 | sense | all | all |
| Class | piRNA | piRNA | 2 | 5'Anchored | sense | all | 18–21 |
| Class | CSR | CSR Class<br>22G | 3 | Partial | antisense | G | 21–23 |
| Class | WAGO | WAGO<br>Class 22G | 3 | Partial | antisense | G | 21–23 |
| Class | ALG | ALG Class<br>26G | 3 | Partial | antisense | G | 26 |
| Class | ERGO | ERGO<br>Class 26G | 3 | Partial | antisense | G | 26 |
| Class | ALG | ALG target<br>22G | 3 | Partial | antisense | G | 21–23 |
| Class | ERGO | ERGO<br>target 22G | 3 | Partial | antisense | G | 21–23 |
| Class | unk | unclassified<br>22G | 3 | Partial | antisense | G | 21–23 |
| Class | unk | unclassified<br>siRNA | 4 | Partial | both | all | 21–24 |

The read counts obtained from tiny-count were used to generate genome-wide coverage profiles for the 22G-RNA and piRNA. The coverage was normalized as reads per million (RPM) and output as BigWig (bw) files using rtracklayer^5^. Differential Expression was computed by using a modified version of tiny.deseq.r, the R module was adjusted to remove the ribosomal RNA (rRNA) from the analysis, and to calculate the gene expression in RPM in addition to DESeq normalized counts. Changes in expression were presented as log2 fold-change, and genes with an adjusted p-value of < 0.05 (after correction with Benjamini & Hochberg)^6^, were considered significant.

#### piRNA target expression

Predicted piRNA targets and CLASH-identified targets of *hsp-70* and *F44E5.4* (stringent targeting rules in *C. elegans*) were obtained from piRTarBase^7^. Mean expression in RPM of the corresponding target piRNA was calculated for each condition.

#### mRNA seq analysis

RNA-seq samples were analyzed and processed using the nf-core/rnaseq (v3.12.0) pipeline, which was executed with Nextflow^8^ (v22.10.6). Sequence quality was assessed with FastQC^9^ (v0.11.9), and low-quality reads, and adapters were removed using Trimmomatic^10^ (v0.67). The processed reads were then aligned to the *C. elegans* genome (WBcel235, Ensembl release 111) with STAR^11^ (v2.7.9a) using default settings. Alignments were quantified against the WBcel235 annotation with Salmon (v1.9.0). Alignment quality control was performed with Qualimap^12^ (v2.2.2) and RSeQC^13^ (v3.0.1).

#### Differential Expression

Differential expression analysis was performed between RNA-seq samples using DESeq2 with the Wald test. Expression changes were presented as log2 fold-change. Genes with an adjusted p-value < 0.05, following correction with the Benjamini & Hochberg^6^ method, were considered significantly differentially expressed.

#### Analysis of Alternative Splicing Events

To identify differential alternative splicing events, RNA-seq data were re-processed using the nf-core/rnasplice (v1.0.4) workflow. This pipeline was executed with Nextflow^8^ (v23.10.0) and relied on reproducible software environments from Bioconda and Biocontainers.

Within the rnasplice pipeline two modules were used to detect differential splicing and calculate the Percent Spliced-In (PSI) for each event, SUPPA2^14^ (v2.3) and rMATS (v4.1.2). Splicing events were considered significant if the absolute difference in PSI (|ΔPSI|) between the 30 min heat shock and control conditions had a p-value < 0.1.

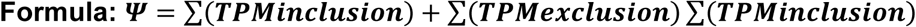

Pearson’s Chi-squared test with Yates’ correction was used to determine if the number of up- or down-regulated splicing events during heat shock differed between the WT and *prde-1* strains (p-value < 0.05).

Differential Transcript Usage (DTU) analysis was performed using SUPPA2^14^. First, the usage of transcripts (isoforms) was tested for significant differences between conditions. Isoforms with a p-value < 0.1 were considered significant. For analysis and visualization, the average Percent Spliced In (PSI or Ψ) value for each isoform was calculated across the biological replicates within each condition. The resulting list of PSI per isoforms was then filtered to retain only those occurring in genes previously identified as differentially expressed (DEGs). Student t-test was used to compare the means between the conditions at significance level p-value < 0.05.

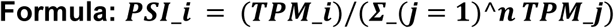

### LONG-READ RNA SEQUENCING

#### Experimental design

Alternative splicing–mediated NMD (AS-NMD) was assessed using the strains and heat-shock conditions described above and the RNAi conditions described below. Briefly, WT and *prde-1 (-/-)* animals were profiled under two RNAi conditions—empty vector control (L4440; NMD on) and *smg-2* RNAi (NMD off)—and two temperature treatments (control and 30 min heat shock at 34 °C). Each strain × RNAi × temperature combination was profiled in three biological replicates (n = 3; 24 libraries total).

#### Library preparation and sequencing

Barcoded *C. elegans* cDNA libraries (WT and *prde-1*) were prepared using 12-plex Kinnex cDNA kits and sequenced across two PacBio Revio SMRT cells. Each SMRT cell multiplexed two libraries: lib01 (Kinnex adapter bcM0001) and lib02 (adapter bcM0004). Genotypes were balanced across barcodes within each library to minimize batch effects.

#### Read processing and alignment

HiFi reads were processed using the Kinnex/Iso-Seq workflow (https://github.com/PacificBiosciences/IsoSeq). Kinnex adapters were identified and split with Skera (v1.40), and cDNA barcodes were demultiplexed with Lima (v2.9.0). Following artificial concatemer removal, full-length non-chimeric (FLNC) reads were generated using isoseq refine. Per-sample FLNC BAM files from both SMRT cells were merged using pbmerge. Merged BAMs were aligned to the *C. elegans* WBcel235 genome (Ensembl release 111) with pbmm2 (ISOSEQ preset) and indexed using samtools. Transcript identifiers were mapped from WormBase gene IDs to gene symbols using a custom transcript-to-gene mapping table.

#### Transcript and gene quantification

Long-read expression and structural categories were quantified with IsoQuant^15^ (v3.13.1) in reference-based mode using the WBcel235 genome and gene annotation. All 24 aligned BAM files were processed simultaneously with the --data_type pacbio_ccs and --fl_data flags to extract per-sample counts. Custom R scripts were used to export gene- and transcript-level count and transcripts per million (TPM) matrices for downstream analysis. SQANTI3^16^ (v6.0.1) was run for isoform quality control using the SQANTI3 QC and SQANTI3 filter modules with default parameters.

#### Alternative splicing–mediated NMD (AS-NMD) benchmarking

To identify high-confidence AS-NMD targets from long-read data, alternative splicing changes were evaluated by comparing L4440 (NMD on) with *smg-2* RNAi (NMD off) across strains (WT, *prde-1*) and temperature conditions (control, 30 min heat shock). Four complementary bioinformatics approaches were applied to the IsoQuant expression outputs:

##### Differential transcript usage (DTU)

DRIMSeq^17^ (v1.38.0) was used to evaluate DTU. Genes and isoforms were filtered to require ≥10 counts across all samples and within at least one genotype group. A gene-level model (∼ group) was fit, and the *smg-2* coefficient was tested via dmTest. Significance was defined at a Benjamini–Hochberg adjusted P < 0.05. Isoform-level confirmation within significant genes was performed using stageR (α = 0.05).

##### Differential exon usage (DEU)

DEXSeq^18^ (v1.56.0) was used to evaluate DEU. Datasets were modeled with the design ∼ sample + exon + condition:exon. Significant DEU was defined at an adjusted gene-level P < 0.05, and selected genes (intersection of genes present across the four contrasts) were visualized using plotDEXSeq.

##### Isoform switching

Isoform switching was analyzed with IsoformSwitchAnalyzeR^19^ using a filtered GTF subset matching quantified transcripts. Splicing switches between L4440 and smg-2 were tested via isoformSwitchTestDEXSeq. Significant switches required a false discovery rate (FDR) q < 0.05 and a minimum isoform fraction difference |ΔIF| ≥ 0.1.

##### Event-level splicing

Annotated alternative splicing events were quantified with SUPPA2^14^ (v2.4) using IsoQuant transcript TPMs (distinct from the Illumina rnasplice/SUPPA2 analyses described above). Local event definitions were generated from the *C. elegans* WBcel235.111 GTF (suppa.py generateEvents -f ioe -e SE SS MX RI FL), producing SE, alternative splice sites (SS; reported as A3/A5), MX, RI, and first/last exon (FL; reported as AF/AL) events.

Percent spliced-in (PSI) was calculated with psiPerEvent requiring a minimum total TPM of 1 for transcripts defining each event (--total-filter 1). Differential splicing was tested with diffSplice using the empirical method and gene correction (-m empirical -gc) on three biological replicates per condition group. For AS-NMD analyses, L4440 was compared with smg-2 RNAi within each strain × temperature group (four primary contrasts: WT and *prde-1* × control and 30 min heat shock). ΔPSI was defined as PSI(smg-2) − PSI(L4440) (positive = higher inclusion under NMD inhibition). Events with P < 0.05 and |ΔPSI| > 0 were called significant. SUPPA2 was used as an independent event-level layer and was not included in the primary consensus gene set.

#### Consensus gene calling and validation

For primary consensus AS-NMD gene identification, significant gene lists from DRIMSeq/stageR, DEXSeq, and IsoformSwitchAnalyzeR were merged per contrast. SUPPA2 was retained as an independent, event-level cross-validation layer and was excluded from the primary consensus calculation.

To profile structural features within these targets, we established a consensus union (p. adj < 0.05 or q < 0.05 across the three primary methods) and extracted IsoQuant assignment events across replicates. Alignment artifacts and poly(A)-site events were filtered out. Structural assignment features were simplified: incomplete splice matches (ISM), full splice matches (FSM), mono-exonic reads, and transcription start/end sites (TSS/TES) were collapsed into a single “consistent” category. For each sample, the absolute number of unique genes containing ≥1 assigned read for a given assignment was calculated.

#### Splice logos

Sequence logos were generated for alternative 3′ splice-site (A3) events called by SUPPA2 from long-read data. Per contrast, A3 events meeting the significance criteria (|ΔPSI| > 0 and P < 0.05) were retained. Each event was parsed into donor–acceptor junction coordinates. Genomic sequence windows were extracted from the *C. elegans* WBcel235 genome using IsoQuant-derived junction coordinates and strand-aware retrieval so that minus-strand sequences were reverse-complemented to transcript orientation. Motif windows were defined as: 5′ splice sites, 9 nt (3 exonic + 6 intronic bases; canonical GT at positions 4–5); 3′ splice sites, 7 nt (6 intronic + 1 exonic base; canonical AG at positions 5–6). Logos were constructed from unique junctions (one sequence per distinct chrom/donor/acceptor/strand, unweighted by read support) with ggseqlogo^20^.

#### Long-read visualization of shared A3 events

Sashimi plots were generated using ggsashimi^22^ and Integrative Genomics Viewer (IGV)^23^. Genes with significant A3 events in all four AS-NMD contrasts (n = 19) were selected for generating the A3 heatmap and for sashimi plots.

### GENE ONTOLOGY

Gene Ontology analysis was performed by using Over Representation Analysis (ORA), with the R package clusterProfiler^24,25^. GO annotations for *C. elegans* were obtained from R package org.Ce.eg.db: Genome wide annotation for Worm^26^ (version 3.18.0). Ontology terms enriched with an adjusted p-value <0.05 or <0.1 (Benjamini & Hochberg)^6^ were considered significant. Fold-Enrichment was calculated as the ratio of the frequency of input genes annotated in a GO term to the frequency of all genes annotated to that term.

### DATA VISUALIZATION

The metaprofiles were generated using deepTools^27^ (v3.5.6) by normalizing the coverage by RPM per bp and then modified to plot them using R. Venn diagrams were generated using the eulerr^28^ package (v.7.0.2) in R. Sashimi plots were generated with ggsashimi^22^ and IGV^23^, A3 sequence logos used ggseqlogo^20^ (see Long-read RNA sequencing). Additional plots were generated using ggplot2^29^ (v.3.5.1).

### RNA ISOLATION AND QUANTIFICATION OF RNA LEVELS BY RT-qPCR

RNA was extracted from 30–40 day-one adults exposed to the indicated heat shock durations and recovery times. Animals were collected in 50 μl Trizol and immediately snap-frozen in liquid nitrogen. Samples were thawed on ice, mixed with 200 μl Trizol, and lysed using Precellys 24 homogenizer. RNA was isolated from lysates with chloroform and precipitated with isopropanol and glycogen. After washing with 70% EtOH, RNA pellet was dissolved in nuclease-free water and treated with TURBO DNase. cDNA was synthesized using iScript cDNA synthesis kit (Bio-Rad; Cat #1708891). qPCR was performed using PowerUp SYBR Green Master Mix (Applied Biosystems; Cat #A25742) on a QuantStudio 3 Real-time PCR System (10 μl reaction, 96-well plate).

The amounts of RNA were quantified using one of the following methods:

#### mRNA RT-qPCR

i. <u>Relative quantification using the ΔΔCt method:</u> mRNA (exon-exon junction primers), and pre-mRNA (intron-exon primers) levels were computed relative to *pmp-3* (previously verified by use to remain steady during heat shock)^1^, which served as an internal control. For NMD-related gene expression analysis, *eft-3* was used as internal control, as previously described^30^. Primer locations are depicted in Method Figure 1 below and primer sequences are in Method Table 2.
ii. <u>Absolute quantification using standard curves:</u> Absolute amounts of pre-mRNA and mRNA were estimated through the generation of standard curves using serial dilutions of defined concentrations of *hsp-70 (C12C8.1)* and *F44E5.4/.5* mRNA, pre-mRNA, as well as *pmp-3* mRNA. The mRNA and pre-mRNA used to generate each standard curve were obtained by PCR amplification from cDNA or genomic DNA. The primers for the standard curve bound externally to the PCR product regions used in the experiments. The amount of PCR amplicons was quantified using a Qubit 4 fluorometer following gel purification. qPCR was conducted using 10-fold serial dilutions of purified PCR products. The standard curves were created from the CT values obtained from each dilution (see Method Figure 1).

The amount of mRNA and pre-mRNA were extrapolated from these curves using the formula.

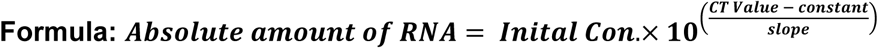

To account for differences in lysis efficiency, variations in harvesting of animals, and other factors, a similar curve was calculated for *pmp-3* and the final values were normalized to *pmp-3* values known to remain stable through heat shock^1^.

**Method Figure 1.**
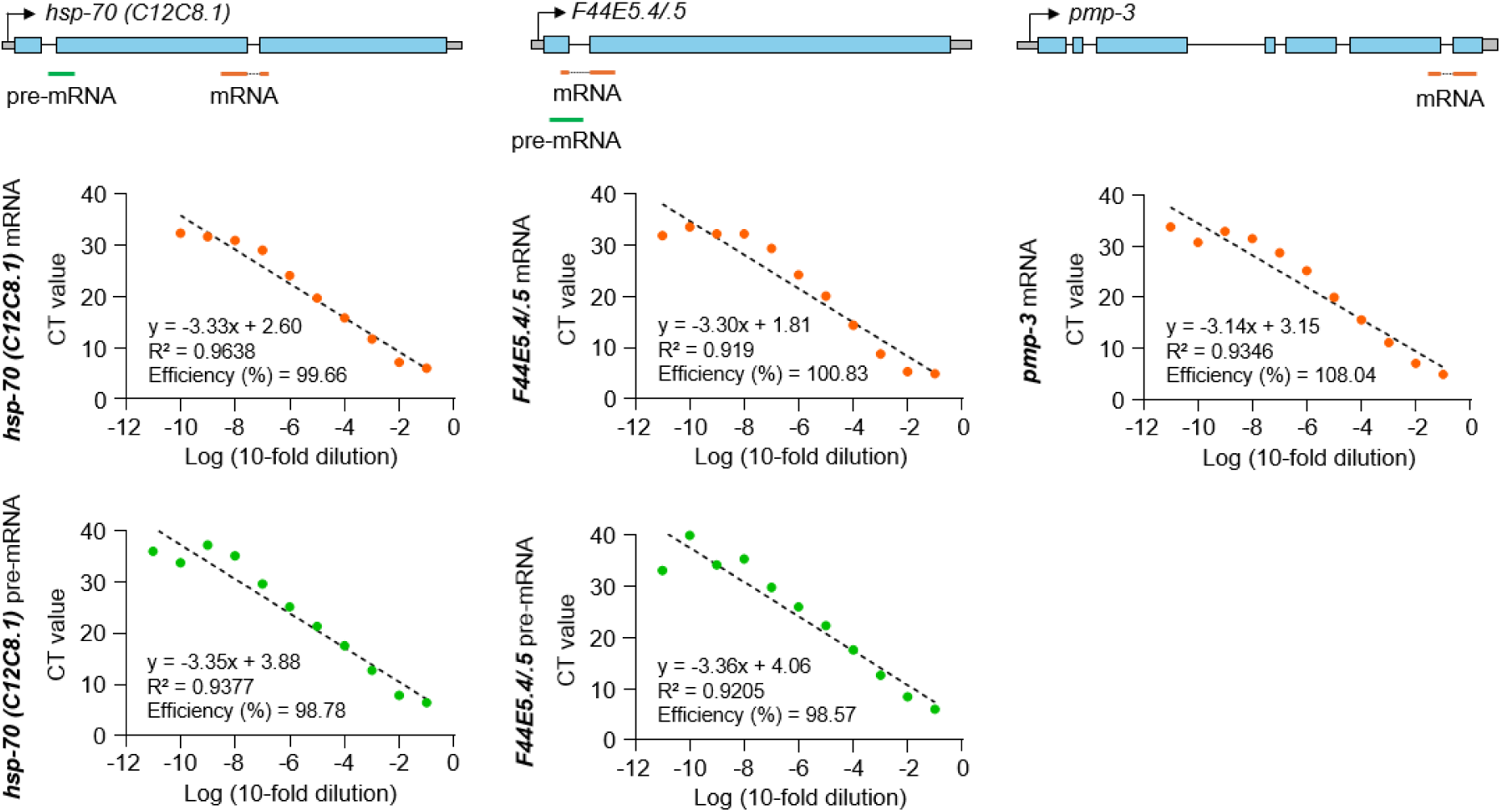
Standard Curves for hsp-70s mRNA and pre-mRNA and pmp-3 mRNA

The primers used for qPCR analysis were designed using Primer 3 software and generated by Integrated DNA Technologies (See Method Table 2).

#### SMALL RNA (sRNA) RT-qPCR

sRNA qPCR was performed by modifying previous stem-loop RT-qPCR protocols^32^. For 22G-RNA detection, total RNA extracted from whole animals or immunoprecipitated RNA was incubated with 50 nM 22G-RNA-specific stem-loop primer at 70°C for 5 min and cooled on ice for ≥ 5 mins. cDNA was synthesized using 10 U/μl Superscript II Reverse Transcriptase (Invitrogen; Cat #18064014) in 1x reaction buffer containing 1 U/μl SUPERase-In RNase inhibitor (RNI; Invitrogen; Cat #AM2696) and 0.5 mM dNTPs. For piRNA detection, total RNA was ligated to 10 μM Universal miRNA cloning linker (NEB; Cat #S1315S) using 10 U/μl T4 Rnl2tr KQ (NEB; Cat #M0373L) at 25°C for 2 hr. Ligated RNA was cleaned up using the Zymo RNA Clean & Concentrator kit, and cDNA was synthesized with the linker stem-loop primer as described for 22G-RNAs detection. The RT-qPCR mix included 2 μl cDNA, 0.25 μM sRNA-specific forward primer, 0.25 μM universal reverse primer, 1x PowerUp SYBR Green Master Mix. Relative fold change of sRNA was quantified relative to *pmp-3* in total RNA or immunoprecipitated RNA using the ΔΔCt method. Primer specificity was tested using a synthetic sRNA (21ur-2348). The specificity of the reaction was verified by running the reaction on agarose gels. Representative gel and PCR design are shown in Method Figure 2. The primer sequences are listed in Method Table 3.

**Table 3.**
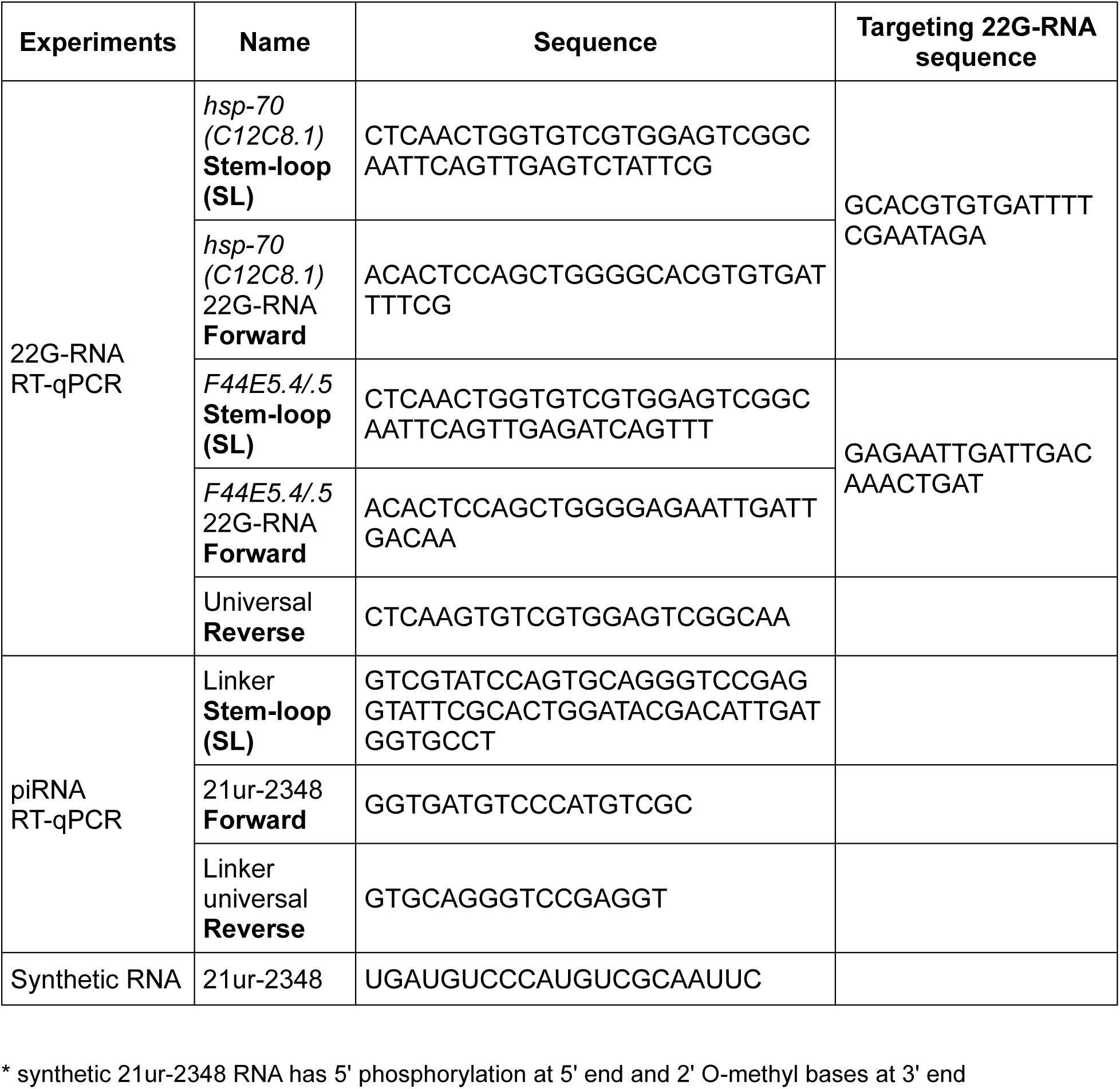
Primer information (small RNA)

**Method Figure 2.**
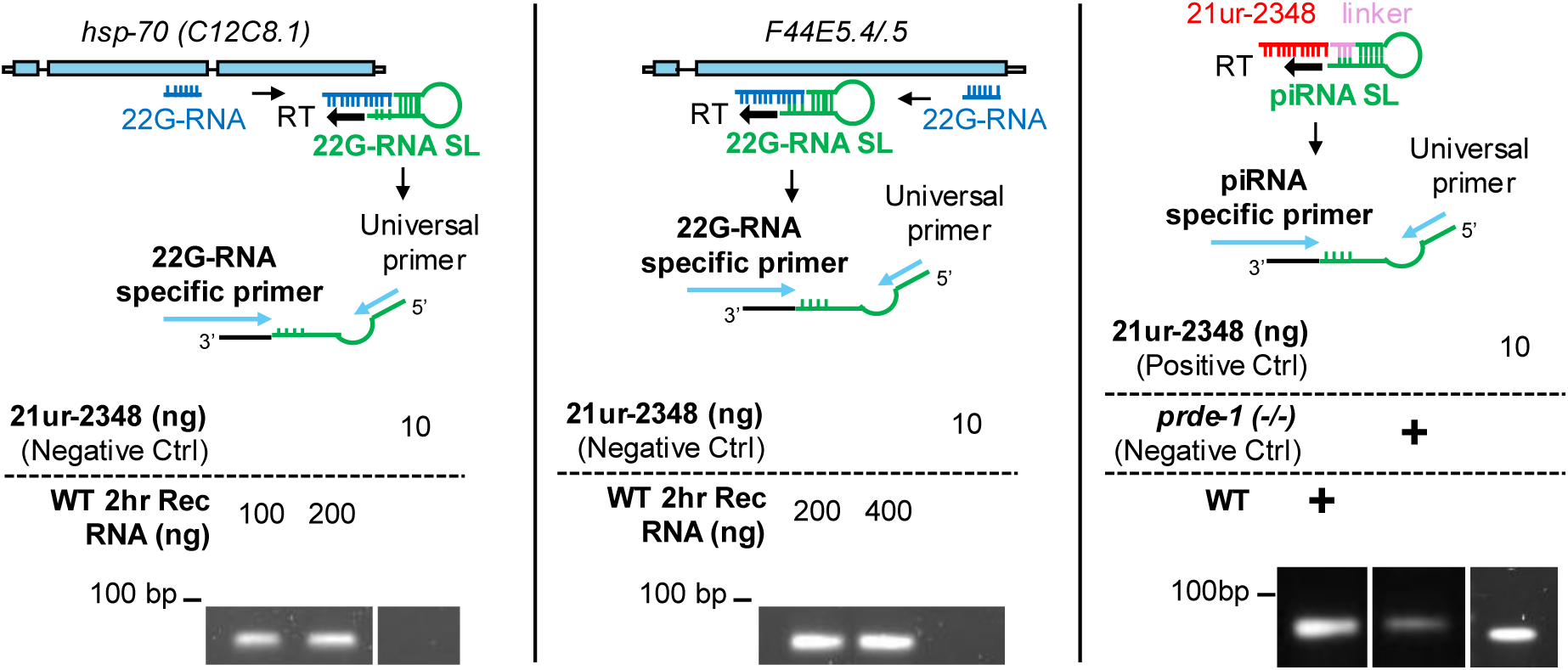
sRNA primers specificity test (expected PCR product size: 69 bp (22G-RNA), 74 bp (21ur-2348))

### EU LABELLING OF NASCENT RNA

To examine the expression patterns of nascent *hsp* genes, we used a pulse-chase approach using the Click-iT Nascent RNA Capture Kit (Invitrogen; Cat #C10365) according to the manufacturer’s instructions. In brief, 50 day-one adult worms were placed on an OP50 bacterial lawn containing 4 mM EU and incubated at 20°C for 10 mins to preload them with EU. Subsequently, the worms were incubated at 20°C (serving as controls) or underwent heat shock for 30 mins at 34⁰C. Control and heat-shocked animals were harvested in 50 µl TRIzol and immediately snap-frozen in liquid nitrogen. To assess maturation of pre-mRNA to mRNA, animals were allowed to recover in the absence of EU. For this, worms were transferred, immediately after heat shock, from EU-containing OP50 plates onto new NGM plates seeded with regular OP50. The plates were then incubated at 20°C for 2, 4, or 6 hours, and subsequently harvested in Trizol. RNA isolation was performed as described in the RT-qPCR protocol. EU-labeled RNA was biotinylated with 0.4 mM biotin azide and purified by reprecipitating. The biotinylated RNA was captured using 12 µl of streptavidin beads. cDNA synthesis was carried out directly on the beads containing nascent RNA using the iScript cDNA synthesis kit. qPCR was processed as described in the RT-qPCR protocol. To quantify mRNA and pre-mRNA, absolute amounts of RNA were estimated from standard curves. To assess maturation from pre-mRNA to mRNA, mRNA expression level at each time point was divided by pre-mRNA expression at 30min HS. In both analyses, mRNA and pre-mRNA expression amounts were quantified relative to EU labeled *pmp-3* amounts.

### 4sU TREATMENT

4sU was delivered as a liquid suspension mixed with bacterial food. The bacterial food, OP50, was heat-killed to minimize its metabolism of 4-Thiouridine (4sU). To most optimally deliver 4sU within a short time, about 700 day-one adult worms were rapidly collected in nuclease-free water and then resuspended in S-basal containing 10 mg/ml heat-killed OP50 and 4mM 4sU (Sigma Aldrich; Cat #T4509). The suspension was seeded onto empty NGM plates to prevent hypoxia (a critical step since *C. elegans* suspended in liquid media have a markedly different heat shock response). The worms, bacteria and 4sU were allowed to dry at RT for 1 hr, and subjected to one of three conditions [not heat shocked (NHS), heat shocked at 34⁰C for 30 mins (30min HS), or heat shocked at 34⁰C for 30 mins and allowed to recover for 2 h at 20⁰C (30min HS + 2hr Rec)], and then crosslinked by exposure to UV (365 nm) for 30 sec. Samples were harvested in nuclease-free water and snap-frozen in liquid nitrogen. 4sU treatment was confirmed not to affect the expression of *hsps* by comparing HSP mRNA levels in the presence and the absence of 4sU. RT-qPCR assay and analysis were performed as described above.

Relative mRNA level shown in Method Figure 3 (Y-axis: Relative mRNA. X-axis: Treatment. Bars: mean value. Error bars: standard error. *n* = 3 biologically independent experiments. Statistical analysis: two-tailed unpaired t-test.).

**Method Figure 3.**
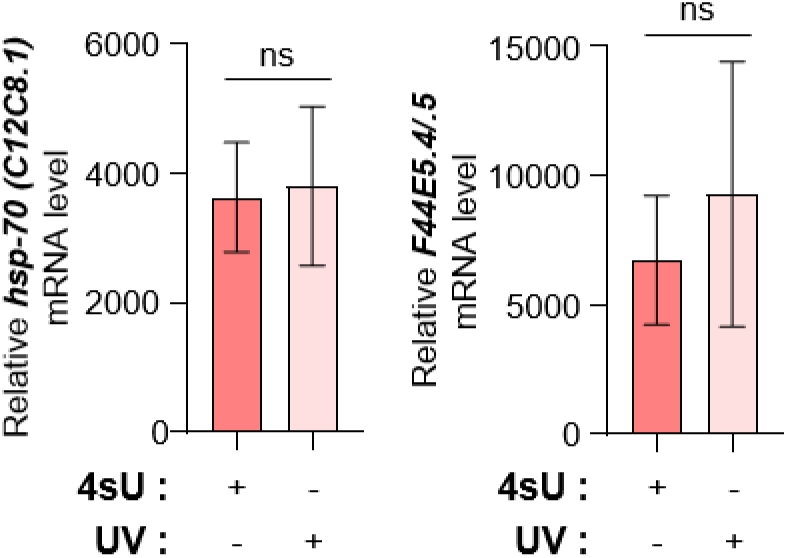
4sU treatment does not impair hsp-70s expression

### RNA IMMUNOPRECIPITATION (RIP) FOLLOWED BY RT-qPCR

For native RIP assay, approximately 700 day-one adults per condition, synchronized by bleach hatching, were harvested in nuclease-free water and snap-frozen in liquid nitrogen. Native or 4sU treated samples (described in the 4sU treatment) were washed once with ice-cold FA buffer (50 mM HEPES/KOH (pH 7.4), 150 mM NaCl, 50 mM EDTA, 1% Triton X-100, 0.1% sodium deoxycholate, 0.5% sarkosyl) and rapidly resuspended in 200 μl FA buffer freshly supplemented with 1 mM DTT, 1x protease inhibitor cocktail (PIC; Thermo Scientific; Cat #78429), and 0.1U/μl RNI. The high detergent content was necessary to minimize non-specific binding, and the high EDTA concentration was required to inhibit Mg²⁺/Ca²⁺-dependent nuclease activity, which we have found is extraordinarily high in *C.elegans* lysates. Worms were lysed using a Precellys 24 homogenizer and sonicated (Bioruptor Pico; 3 cycles of 30 s on/off). Debris was removed by centrifugation (13,500 rpm at 4°C for 5 mins). For PRG-1, 4sU-treated RNP-6, and 4sU-treated total Pol II RIP experiments, Protein A/G Magnetic Beads (20 μl per sample; Thermo Scientific; Cat #88803) were used. Beads were washed twice with FA buffer and 5 μl of beads were added to the lysates and incubated at 4°C for 1 hr to eliminate non-specific binding. 10% of the pre-cleared lysate was kept as input, and the rest was incubated with anti-FLAG M2 (Sigma-Aldrich, Cat #F3165-1MG), anti-HA (C29F4) (Cell Signaling Technology; Cat #3724S), anti-total Pol II (Diagenode; Cat #C1520004) for 2–3 hr at 4°C, followed by 1 hr incubation with 15 μl beads. Beads were washed sequentially with low salt, high salt, LiCl, and TE buffers. For MOG-7 RIP experiments, GFP-Trap Magnetic Particles M-270 Kit (ChromoTek; Cat #gtdk-20) was used by modifying the manufacturer’s instructions. In brief, beads were washed with the manufacturer’s washing buffer containing 0.005% Tween-20 for 15 min at RT, washed three times with FA buffer, and pre-blocked with salmon sperm DNA (Invitrogen; Cat #15632011) for 1 hr at 4°C. After aliquoting 10% of lysate as input, the remaining lysate was incubated for 2–3 hr at 4°C with pre-blocked GFP beads. Beads were washed sequentially with the washing buffer, and TE buffer. After washing, beads were eluted with 150 μl elution buffer (10 mM EDTA, 1% SDS, 0.1 M sodium bicarbonate) containing 0.1U/μl RNI. Elutes and volume-matched inputs were de-crosslinked by incubating them with 0.8U proteinase K (NEB; Cat #P8107S) for 1 hr at 45°C. RNA was isolated using Trizol and mRNA and sRNA RT-qPCR was performed as described above. RT-qPCR analysis was conducted differently across the experiments as following:

i. PRG-1 experiments Because we had no reason to expect that *pmp-3* would also bind to PRG-1, and be immunoprecipitated, to control for variation in harvesting, lysis, loading etc., we normalized *hsps* in the IP sample to *pmp-3* in the input sample using the ΔΔCt method. *bath-45*, known to interact with PRG-1 was used as a positive control for the efficacy of RIP. WT animals not expressing FLAG were used as a negative control to account for the non-specific interactions of the abundant *hsp* mRNA generated by heat shock.
ii. Total Pol II experiments Absolute amounts of mRNA and pre-mRNA were estimated using standard curves and normalized to immunoprecipitated *pmp-3*. The 22G-RNA in the IP sample was also normalized by *pmp-3* in the same IP sample.
iii. RNP-6 and MOG-7 experiments *hsps* pre-mRNA and 22G-RNA in the IP sample were normalized to *pmp-3* in the input sample using the ΔΔCt method (same reason as in (i)). WT animals not expressing HA and GFP were used as a negative control to account for the non-specific interactions of the abundant *hsp* mRNA generated by heat shock.

The sequence of the primers used for RIP experiments are listed in Method Table 2 and Method Table 3.

### CHROMATIN IMMUNOPRECIPITATION (ChIP) FOLLOWED BY qPCR

Chromatin immunoprecipitation (ChIP) was performed as previously described^30^. Following exposure to indicated conditions, approximately 700 day-one adults, synchronized by bleach hatching, were harvested in 1x PBS and crosslinked with 2% formaldehyde at RT for 10 mins. Crosslinking was quenched with 250 mM Tris (pH 7.4) for 10 mins. Samples were washed three times in ice-cold 1× PBS containing 0.1x PIC and then snap-frozen in liquid nitrogen. Pellets were resuspended in 200 μl FA buffer freshly supplemented with 1 mM DTT and 1x PIC. Worms were lysed and sonicated (8-10 cycles of 30 s on/off). Debris was removed by centrifugation (13,500 rpm at 4°C for 5 mins). For HSF-1 ChIP experiments, endogenous HSF-1 tagged with FLAG was immunoprecipitated using anti-FLAG M2 magnetic beads (Sigma-Aldrich; Cat #M-8823). Beads (20 μl per sample) were washed with FA buffer and pre-blocked with salmon sperm DNA. After aliquoting 10% of lysate as input, the remaining lysate was incubated overnight at 4°C with pre-blocked FLAG beads. For total Pol II ChIP experiment, Protein A/G Magnetic Beads (20 μl per sample) were used. Beads were washed with FA buffer and 5 μl of beads were added to the lysates and incubated at 4°C for 1 hr to eliminate non-specific binding. 10% of the pre-cleared lysate was kept as input, and the rest was incubated overnight at 4°C with anti-total Pol II, followed by 3 h incubation with 15 μl beads. Beads were washed sequentially with low salt, high salt, LiCl, and TE buffers, then eluted in 150 μl elution buffer. Elutes and volume-matched inputs were de-crosslinked overnight with 0.8U proteinase K (NEB; Cat #P8107S) and 160 mM NaCl. DNA was purified using phenol/chloroform/isoamyl alcohol. qPCR was performed as above. HSF-1 experiments were expressed as fold change relative to the non-heat shocked control of each genotype, and total Pol II experiments were expressed as % input value. The sequence of the primers used for ChIP experiments are listed in Method Table 2 and depicted in Figure 2c and Supplementary Figure 1f.

### SUBCELLULAR FRACTIONATION

Subcellular fractionation was performed with ∼700 day-one adults per condition synchronized through bleach hatching method. Worms were harvested in 1.5 ml tube with nuclease-free water, pelleted and snap-frozen in liquid nitrogen. Frozen pellet was washed with 500 μl hypotonic buffer (15 mM HEPES/KOH (pH7.5), 10 mM KCl, 5 mM MgCl2, 0.1 mM EDTA, 350 mM Sucrose). Worms were resuspended in 100 μl hypotonic buffer supplemented with 1 mM DTT, 1x PIC and 0.1U/μl RNI and homogenized by pestle (SP Bel-Art; Cat #F19923-0001). To remove worms’ bodies and debris, homogenates were centrifuged at 500 xg at 4°C for 5 mins, and supernatant was transferred to a new 1.5 ml tube. The supernatant was then centrifuged at 4,000 xg at 4°C for 5 mins to collect the nuclear (pellet) fraction. The nuclei pellet was washed three times with 200 μl hypotonic buffer. After last washing, the pellet was resuspended in 120 μl nucleus lysis buffer (50 mM Tris (pH7.5), 140 mM NaCl, 1.5 mM MgCl2, 0.5% (w/v) NP-40) supplemented with 1 mM DTT, 1x PIC and 0.1U/μl RNI, and incubated in ice for 20 mins. 20 μl of sample was aliquoted to a 1.5 ml tube labelled as “input”, and the remaining samples were centrifuged at 20,000 xg at 4°C for 10 mins. The supernatant was transferred to a 1.5 ml tube labelled as “nucleoplasm”. The pellet was resuspended in 100 μl high salt solution (10 mM Tris (pH7.5), 700 mM NaCl) supplemented with 1 mM DTT, 1x PIC and 0.1U/μl RNI, and incubated in ice for 20 mins, and centrifuged at 20,000 xg at 4°C for 5 mins. The supernatant was transferred to a 1.5 ml tube labelled as “Chromatin”. The RNA in each sample was isolated using Trizol, and *hsp* gene expression level was measured as described in the RT-qPCR protocol. The results of experiments were presented as percentage input values.

This method of extraction did result in the loss of some nuclei. Nevertheless, the nuclei obtained were ‘clean’ and not contaminated with cytoplasmic fractions, confirmed by conducting a Western blot and probing for tubulin.

### SINGLE MOLECULE FLUORESCENT IN SITU HYBRIDIZATION (smFISH)

smFISH was performed as previously described^30^. Probes were designed against mRNA according to the Biosearch Technologies website, and consisted of 11 different 25–35 bp sequences that were complementary to the *hsp-70s* (*C12C8.1* and *F44E5.4/.5*) mRNA. About 20 day-one adult worms were harvested in 1x PBS and fixed in 4% paraformaldehyde (Electron Microscopy Sciences; Cat #50-980-487) for 45 mins. After two washes with 1x PBS, worms were permeabilized in 70% ethanol at 4°C for 24–26 hr. Worms were washed with Stellaris Buffer A (Biosearch Technologies; Cat # SMF-WA1-60) and incubated at 37°C for 16 hr in hybridization buffer (Biosearch Technologies; Cat #SMF-HB1-10) containing 0.2 mM FISH probes targeting *hsp-70s* (*C12C8.1* and *F44E5.4/.5*) mRNA. After washing three times with pre-warmed Wash Buffer A and one time with Wash Buffer B (Biosearch Technologies; Cat # SMF-WB1-20), worms were mounted in Vectashield with DAPI (Vector Laboratories; Cat #H-1200-10). Imaging was performed using a Leica TCS SPE Confocal Microscope with 20x dry and 63x oil objectives, and LAS AF software.

### IMMUNOSTAINING

Immunostaining of dissected gonad was performed as previously described^30^. 10–15 day-one adults were picked into 15 μl 1x PBS on a cover glass (Leica Biosystems; Cat #3800105) and dissected with a surgical blade #11. Dissected worms were fixed in 4% paraformaldehyde for 6 mins. A charged slide (Epredia; Cat #4951PLUS-001) was placed over the cover glass, then quickly transferred onto a metal block that had been cooled with dry ice for over 10 min. After quickly removing the cover glass, samples were post-fixed in 100% methanol pre-chilled at −20°C for 2 min, washed in 1x PBST (1x PBS with 0.1% Tween-20), and then blocked in 1x PBST with 0.5% BSA for 1 hr. Samples were incubated with primary antibody overnight at 4°C, followed by secondary antibody for 2 hr at RT. After washing, they were mounted in the Vectashield with DAPI and imaged. For PRG-1 and HSF-1 immunostaining, Mouse anti-FLAG M2 (1:2000) was used. For PRDE-1 immunostaining, Rabbit anti-HA (1:1000) was used. Donkey anti-Mouse Alexa Fluor 488 (1:500; Invitrogen; Cat #A32766) and Goat anti-Rabbit Alexa Fluor 647 (1:500; Invitrogen; Cat #A21245) were used as secondary antibodies.

### SMALL RNA INJECTION AND RECOVERY ASSAY

Small RNA was prepared from total RNA extracted from

i. WT animals subjected to 30 min HS
ii. *prde-1* animals subjected to 30 min HS

Small RNAs were fractionated using the Mirvana miRNA Isolation Kit. The small RNA was then injected at a concentration of 50 ng/μl. Following injection, worms were recovered at 20°C for 2 hr and subsequently maintained under non-heat-shock conditions (NHS) or subjected to 30 mins heat shock at 34°C. Worms were collected in 1x PBS after 6 hr incubation at 20°C and used for *F44E5.4/.5* mRNA FISH experiment. For RT-qPCR analysis, worms were collected immediately after heat treatment in Trizol, and their RNA then harvested and quantified as described above.

### RNA INTERFERENCE (RNAi)

RNAi experiments were conducted using the standard feeding RNAi methods. Bacterial clones expressing the control (empty vector pL4440) construct and the dsRNA targeting different *C. elegans* genes were obtained from the Ahringer RNAi library^31^, now available through Source Bioscience. All RNAi clones used in experiments were sequenced for verification. For RNAi experiments, RNAi bacteria with empty (pL4440 vector as control) or the dsRNA-expressing plasmid were grown overnight in LB liquid culture containing ampicillin (100 μg/ml) and then induced with IPTG (1 mM) for 2 hr before seeding the bacteria on NGM plates supplemented with ampicillin (100 μg/ml) and IPTG (1 mM). Bacterial lawns were allowed to grow for 2 days before the start of the experiment. RNAi-seeded plates were used for RT-qPCR experiments to measure NMD-related gene expression, to evaluate the hatch rate of embryos, and for immunoblot assays (to analyze ubiquitinylated proteins). For RT-qPCR and Western blot experiments, worms were initially grown on OP50 plates for 24 hours after egg laying or bleaching, respectively. Then, larvae were transferred to RNAi plates and grown for 60–62 hours before harvesting. This was done to bypass the embryonic lethality of *smg-1* or *smg-2* RNAi.

Feeding RNAi with empty vector (L4440) or *smg-2* was also used for PacBio Kinnex long-read AS-NMD experiments (see Long-read RNA sequencing).

### EMBRYO HATCH RATE

Embryo hatch rate was assessed using 2–5 day-one adult worms, which laid eggs for different durations following NHS or 30min HS, as described previously^30^. After removing mothers, embryos were counted and incubated at 20°C. The number of live progenies were scored after 72 hr. For RNAi experiment, mothers were prepared by transferring them to RNAi plates at L4-stage and allowed to lay eggs for 12 hr after treatment.

### WESTERN BLOT

Worm lysates were prepared by boiling samples in Laemmli sample buffer (Bio-Rad; Cat #1610737) with β-mercaptoethanol. For ubiquitinylated protein analysis, about 150 worms were prepared as described in the RNAi protocol. The worms were collected, snap frozen, and lysed by boiling and grinding using a pestle. Lysates were separated on SDS-PAGE (HSP-70: 10%, Ubiquitinylated protein: 12%) and transferred onto nitrocellulose membrane (Bio-Rad; Cat #1620115). Membranes were blocked at RT for 1 hr in Odyssey Blocking Buffer (LI-COR; Cat #927-50000), incubated overnight at 4°C with primary antibodies, and then with IRdye goat anti-mouse IgG 800CW (1:10,000; LI-COR; Cat #926-32210) at RT for 2 hr. Imaging was performed using the LI-COR Odyssey Infrared System and analyzed with Image Studio software.

To detect HSP-70 levels, mouse anti-FLAG M2 (1:1000) was used to detect HSP-70::FLAG in strains where endogenous *hsp-70* (C12C8.1) was tagged with CRISPR/Cas9. Mouse anti-Ubiquitinylated proteins, clone FK2 (1:1000; Sigma-Aldrich; Cat #04-263), was used to detect ubiquitinylated proteins. For tubulin detection as a loading control, mouse anti-tubulin, clone AA4.3 (1:1000; DSHB; Cat #AB_579793) was used.

### ANALYSIS OF mRNA VARIANTS

To assess whether the loss of piRNAs led to increased transcript errors, we devised a method to identify sequences of individual mRNA molecules prior to the possible incorporation of errors due to PCR amplification. RNA extracted from 3 biological repeats of: wild type *C. elegans* at 30 minutes at 34⁰C, and *prde-1* at 30 minutes at 34⁰C were used. mRNAs were tagged with UMIs using the SMART-Seq® Total RNA Pico Input with UMIs (ZapR™ Mammalian). The UMIs were attached to mRNAs during the first strand-cDNA synthesis, prior to PCR amplification.

Subsequently, mRNAs were PCR amplified with 3 cycles of amplification for the initial library PCR and 12 cycles of amplification for the enrichment PCR. Sequencing was performed on the NovaSeq 6000; the number of paired-end reads sequenced were in the range of 50 million-80million, with a read size of 100 bp.

Next, data was pre-processed using a modified version of the RNA with UMIs v1.0.16 pipeline^33^ and a modified version of the GATK^34,35^ Best Practice: RNA-seq Variant Calling Workflow using the tools from gatk (v4.6.1.0) and Picard (v3.3.0).

The complete list of relevant tools, and the custom script used are available on github (https://github.com/Prahlad-Lab/umi_celegans_analysis_consensus) DOI 10.5281/zenodo.17873233.

The fastq.gz files required to run this analysis pipeline are available through the European Nucleotide Archive (ENA) at EMBL-EBI under accession number PRJEB101605 (https://www.ebi.ac.uk/ena/browser/view/PRJEB101605).

FastQC^36^ was used to check the quality of the FASTQ paired-end reads. The FASTQ files were converted to unmapped bam (uBAM) using gatk FastqToSam tool. Unique molecular identifiers (UMI) were extracted using fgbio ExtractUmifromBam. The UMI were extracted using the parameters indicated in the SMART-Seq® Total RNA Pico Input with UMIs (ZapR™ Mammalian) kit, the SMART UMI Adapter from read2 was removed by trimming reads a total of 12 nucleotides: “8 nt UMIs + 3 nt UMI linker + 3 nt”. The quality of the uBAM file was measured again with FastQC^36^

Reads were aligned to genome with STAR^11^ (v 2.7.11b) using the Caenorhabditis elegans genomic reference and annotation (ENSEMBL WBcel235 version 114)^37^. Per-Sample two pass-mapping was enabled using the option -twopassMode Basic. The main output was an unsorted BAM that included unmapped reads, --outSAMtype was set to “BAM Unsorted” and -- outSAMunmapped was set to “Within”.

#### UMI Grouping and Consensus Generation

Following alignment, BAM files were sorted by coordinate with Picard SortSam tool, then merged with unaligned Bam file to get the UMI tags. fgbio GroupReadsByUmi was used to create subgroups (families) based on their position, UMI, and, the parameters --strategy Adjacency --edits 1 and --family-size-histogram were used to correct sequencing errors within UMIs themselves and output the family sizes. Around 70-80% of the reads in all the replicates were Family size =1, as shown in the plot (Method Table 4).

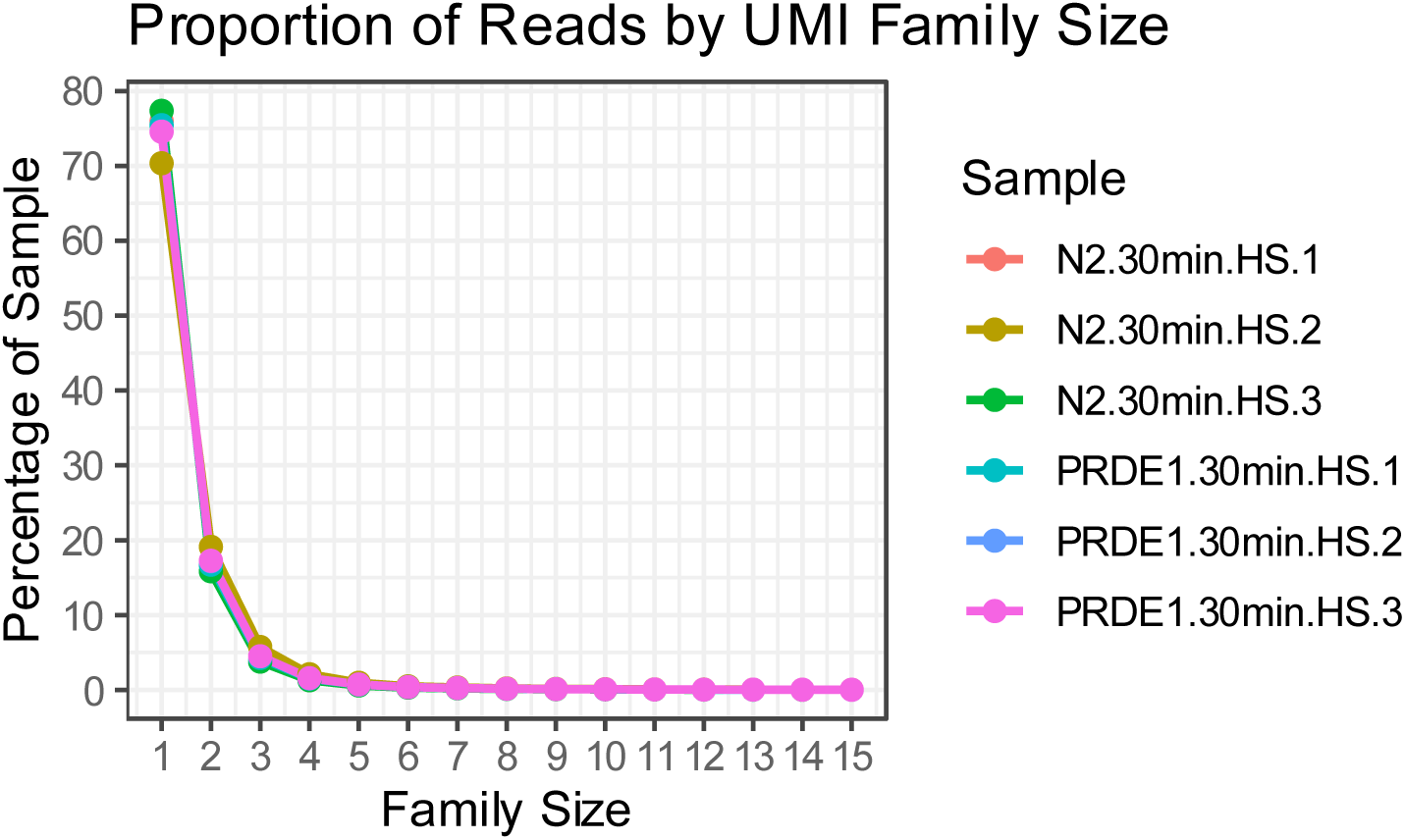

The different families were collapsed into single consensus reads by using fgbio CallMolecularConsensusReads. Families of size 1 were preserved using the parameter --min-reads 1. The consensus reads were mapped again to the WBcel235genome using STAR and merged again to unmapped BAM to get UMI tags.

Following the GATK best practices for RNA-seq Variants, BAM files were processed with gatk SplitNCigarReads to split the reads with “N” in their cigar. To filter out DNA variants, a recalibration model was built with gatk BaseRecalibrator. The resulting model was applied to the consensus BAM files with gatk ApplyBQSR to obtain the BSQR BAM.

#### Pileup Generation and analysis

A pileup of the BSQR BAM was generated with bcftools mpileup using the parameter -d 1000000 to keep the reads at all loci. To increase confidence that we were detecting mRNA variants, and not stochastic nucleotide changes we:

i. analyzed only regions that had generated over 10 reads (i.e. at least 10 mRNAs corresponded to the region to identify the ones which were different). The pileup file was analyzed with a custom python script to filter the regions with total depth <=10 and count the proportion of alternative reads and reference reads.
ii. Excluded regions where “Alternative Read/Total Reads =1.0” or “0.5”. This was used to exclude possible homozygous or heterozygous SNPs with frequencies of 1 or 0.5, respectively, although this does not account for differences in expression from the different alleles.

A custom R script was used to generate tables with the statistics of the Total Number of “mistakes”, the Distribution of the Alternative Reads/Reference reads, a Kolmogorov-Smirnov Test was used to compare the distributions. The approximate mean error rate across all nucleotides (Global error rate) during heat shock was in the order of approximately 10^-3^/bp. GO analysis of the genes with more mismatches in *prde-1* than in WT was performed.

#### Variant Call and Analysis

HaplotypeCaller ^38^ was used with the BSQR BAM to call variants and output VCF file. SnpEff^39^ was used to annotate the vcf with the SnpEff wbcel235 internal annotation. A custom R script was used to read the VCF and filter the variants with total depth ≤ 10, and SnpEFF Functional Impact type of “Modifier”. Variant Sites with a Ratio of Alternative Read/Total Reads = 1.0 were excluded to remove potential genomic Variants. The following were calculated:

i. Transcription error rate: Ratio of Alternative Read/Total Reads
ii. Deleterious Variant Rates: Ratio Deleterious Variants/Total Reads
iii. Mismatch Error rate: change in nucleotides type.
iv. Two sample t-test was used to determine if there was statistical difference between WT and *prde-1 (-/-)* samples, with p-value < 0.05 considered significant.

A total of 1315-1684 nucleotide positions in the approximately 3.7 × 10^7^ individual mRNAs per sample analyzed were identified as high-quality variant-sites. Distribution of the variant across a) different gene size and b) among different nucleotide base changes were tested (Amongst the base changes G>A was the most highly represented).

### STATISTICAL ANALYSIS

Statistical tests, sample sizes, and definitions of replicates are specified in the corresponding Figure Legends. Unless otherwise indicated, tests were two-sided, and statistical significance was defined as P < 0.05; where multiple comparisons were performed, P values were adjusted using the Benjamini–Hochberg false-discovery rate (FDR) correction.

#### RNA-sequencing and small RNA analyses

Wilcoxon test was used to compare the expression of the piRNAs. Enrichment or overlap between gene/RNA sets was assessed by hypergeometric testing using the phyper function in R^40^. Differences in the numbers of up- and down-regulated splicing events between WT and prde-1(-/-) animals were assessed using Pearson’s chi-squared test with Yates’ correction.

Distributions of transcript error rates obtained from pileup analyses were compared using the Kolmogorov–Smirnov test.. All the statistical analyses were done in R^40^. Kolmogorov–Smirnov test was used to compare the probability distributions of mRNA error rates in the pileup analysis.

#### Long-read and alternative-splicing analyses

For long-read SUPPA2 metrics, global differences were assessed with Wilcoxon rank-sum tests and two-way ANOVA (ΔPSI∼strain × event) in R. IsoQuant-derived transcript structural features were compared between WT and *prde-1(-/-)* animals using two-way ANOVA within treatment groups, followed by estimated-marginal-means pairwise comparisons (`emmeans`) with Benjamini–Hochberg FDR correction.

#### qPCR data and phenotypic analysis

Comparisons between two groups were performed using two-tailed unpaired Student’s t-tests; experiments involving multiple factors were analyzed by two-way ANOVA followed by Fisher’s LSD multiple-comparisons test, as specified in the Figure Legends. These analyses were performed using GraphPad Prism. Significance is indicated as: *P < 0.05, *P* < 0.01, and *P < 0.001.

#### Statistical analysis (other)

Unless otherwise indicated, sequencing analyses were performed in R and all additional statistical procedures are described in the corresponding Figure Legends.

## DATA AVAILABILITY

The data for this study have been deposited in the European Nucleotide Archive (ENA) at EMBL-EBI under accession numbers PRJEB101605 (https://www.ebi.ac.uk/ena/browser/view/PRJEB101605), PRJEB101594 (https://www.ebi.ac.uk/ena/browser/view/PRJEB101594), PRJEB101181(https://www.ebi.ac.uk/ena/browser/view/PRJEB101181). https://www.ebi.ac.uk/ena/browser/view/PRJEB101605 https://www.ebi.ac.uk/ena/browser/view/PRJEB101594 (https://www.ebi.ac.uk/ena/browser/view/PRJEB101181) and PRJEB121150 (https://www.ebi.ac.uk/ena/browser/view/PRJEB121150)

---

**Supplementary Figure 1.**
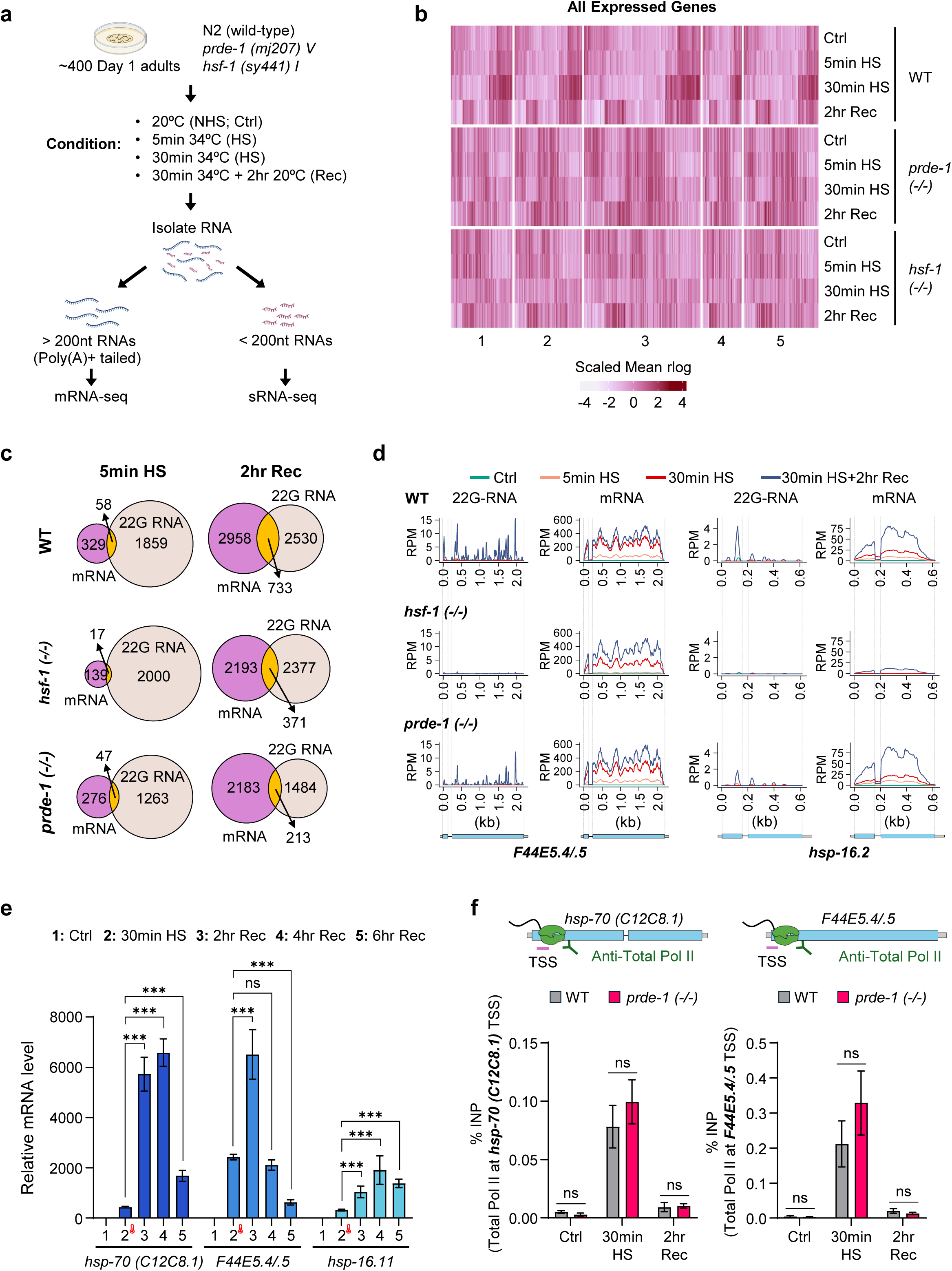
Heat-shock-dependent mRNA and 22G-RNA dynamics in wild-type and piRNA-pathway mutants. **a.** Experimental design for matched mRNA- and small RNA-sequencing. Day 1 wild-type (N2), *prde-1(mj207)*, and *hsf-1(sy441)* adults were maintained under control conditions (Ctrl), exposed to 5 or 30 min heat shock (HS), or allowed to recover for 2 hr after 30 min HS (Rec). Total RNA was fractionated for mRNA- and small RNA-sequencing. **b.** Heat map of mRNA transcript abundance across all expressed genes in wild-type, *prde-1*, and *hsf-1* animals under the indicated conditions. Values represent scaled mean expression. **c.** Overlap between differentially expressed mRNAs and antisense 22G-RNAs after 5 min HS and 2 hr recovery in wild-type, *hsf-1*, and *prde-1* animals. Numbers indicate significantly regulated mRNAs and 22G-RNAs and their overlap. **d.** mRNA and antisense 22G-RNA profiles across *F44E5.4/.5* and *hsp-16.2* in wild-type, *hsf-1*, and *prde-1* animals under control conditions, after 5 or 30 min HS, and after 2 hr recovery. Gene models are shown below. **e.** RT–qPCR analysis of *hsp-70 (C12C8.1)*, *F44E5.4/.5*, and *hsp-16.11* mRNA during heat shock and recovery, showing the kinetics of stress-induced transcript accumulation and clearance. *n* = 3–30 biological repeats. Statistical analysis: unpaired t-test. **f.** RNA polymerase II occupancy at the transcription start sites (TSS) of *hsp-70 (C12C8.1)* (left) and *F44E5.4/.5* (right) in wild-type and *prde-1* animals under control conditions, after 30 min HS, and following 2 hr recovery, measured by total Pol II ChIP–qPCR and expressed as percentage of input. *n* = 3–4 biological repeats **e, f**: Data are mean ± SEM. Statistical analyses: two-tailed unpaired t-test (**e**); two-way ANOVA (**f**). ns, not significant; ✱✱✱ *p* < 0.001.

**Supplementary Figure 2.**
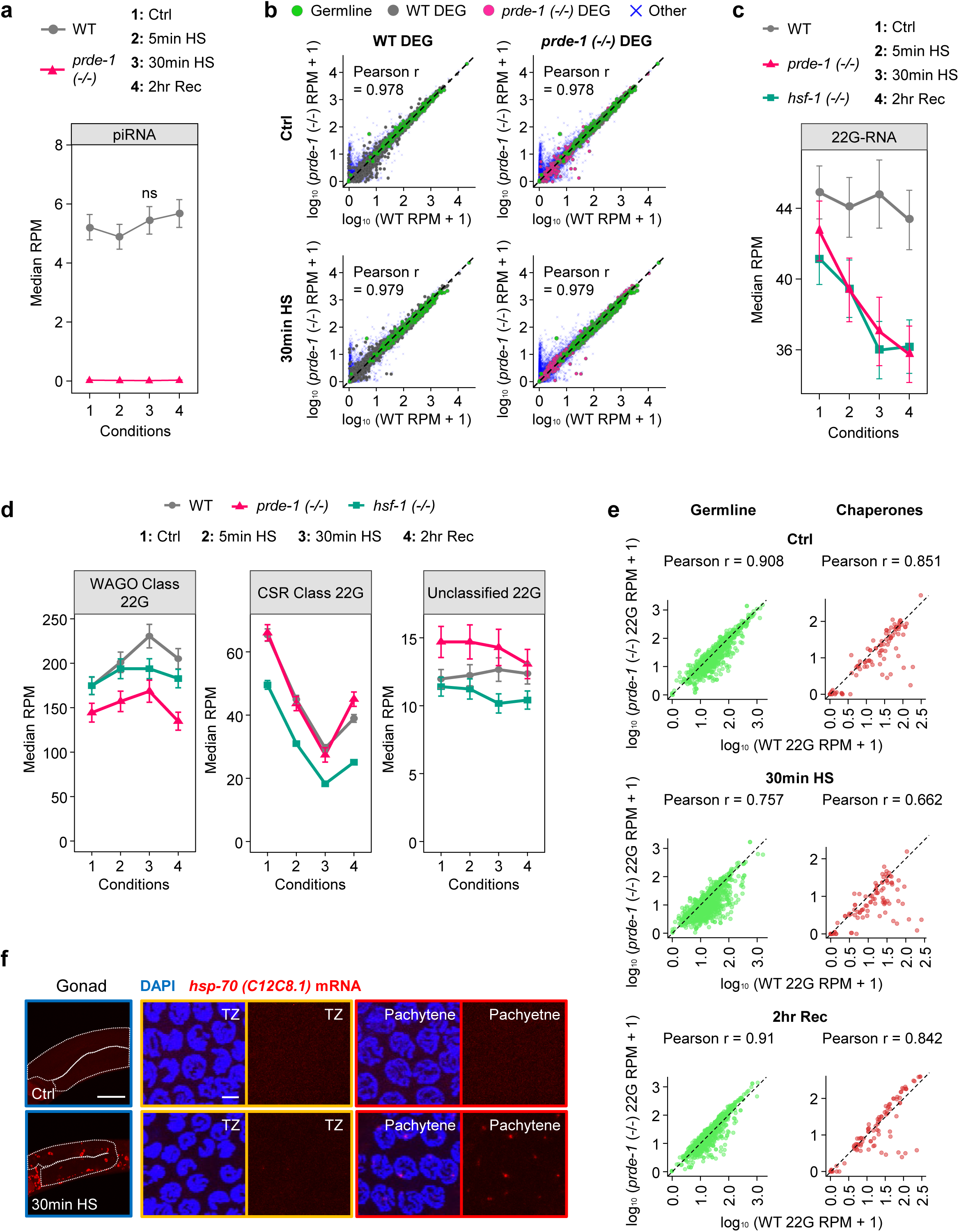
Heat-shock-responsive 22G-RNA production is selectively impaired in *prde-1* mutants. **a.** Median piRNA abundance in wild-type and *prde-1* animals under control conditions, after 5 or 30 min heat shock (HS), and following 2 h recovery (Rec). **b.** Relationship between mRNA abundance in wild-type and *prde-1* animals for wild-type (left) or *prde-1* (right) differentially expressed genes (DEGs), under control conditions and after 30 min HS. Germline-expressed genes are shown in green; DEGs in gray or magenta; other genes in blue. Pearson correlation coefficients are indicated. **c, d.** Global 22G-RNA abundance across genotypes and conditions. **(c)** Median abundance of all 22G-RNAs in wild-type, *prde-1*, and *hsf-1* animals. **(d)** Median abundance of WAGO-class, CSR-1-class, and unclassified 22G-RNAs. **e.** Relationship between 22G-RNA abundance in wild-type and *prde-1* animals for 22G-RNAs targeting germline-expressed genes (left) and molecular chaperone genes (right), under control conditions, after 30 min HS, and following 2 hr recovery. Pearson correlation coefficients are indicated. **f.** RNA FISH showing heat-shock-induced *hsp-70 (C12C8.1)* mRNA (red) in the germline. Schematic indicates the gonadal region imaged; enlargements show the transition zone (TZ) and pachytene regions under control conditions and after 30 min HS. DNA, DAPI (blue). Scale bars, 50 μm (Gonad) and 3 μm (TZ, Pachytene). **a, c, d**: Data are median ± SEM. Statistical analyses: Wilcoxon test (**a**). ns, not significant.

**Supplementary Figure 3.**
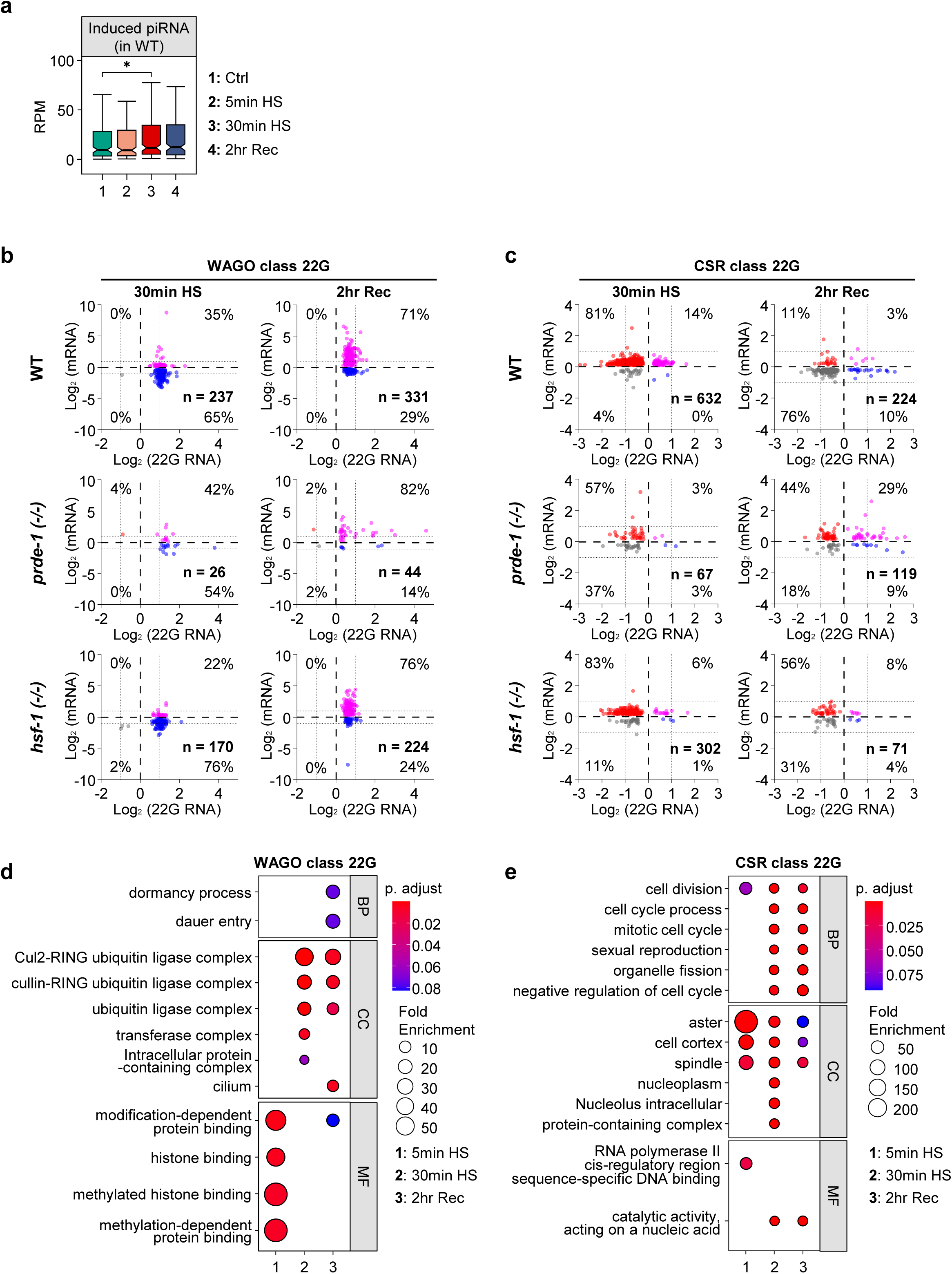
Heat shock induces distinct piRNA- and HSF-1-dependent small-RNA responses. **a.** Abundance of piRNAs induced by heat shock in wild-type animals under control conditions, after 5 or 30 min heat shock (HS), and following 2 h recovery (Rec). Boxplots show the distribution of RPM values for the heat-shock-induced piRNA set. **b, c.** Relationship between changes in 22G-RNA abundance and expression of their cognate mRNA templates for **WAGO-class (b)** and **CSR-1-class (c)** 22G-RNAs after 30 min HS and following 2 hr recovery. Plots show log2 changes in 22G-RNA abundance (x-axis) versus cognate mRNA abundance (y-axis) for wild-type, *prde-1*, and *hsf-1* animals. Dashed lines indicate thresholds used to classify changes; percentages indicate the fraction of transcripts within each quadrant, and *n* indicates the number analyzed. **d, e.** Gene Ontology (GO) enrichment of genes targeted by heat-shock-responsive **WAGO-class (d)** and **CSR-1-class (e)** 22G-RNAs at 5 min HS, 30 min HS, and 2 hr recovery. Biological process (BP), cellular component (CC), and molecular function (MF) terms are shown. Circle size indicates fold enrichment and color indicates adjusted *p* value. **a**: Data are median ± SEM. Statistical analyses: Wilcoxon test. ✱ *p* < 0.05

**Supplementary Figure 4.**
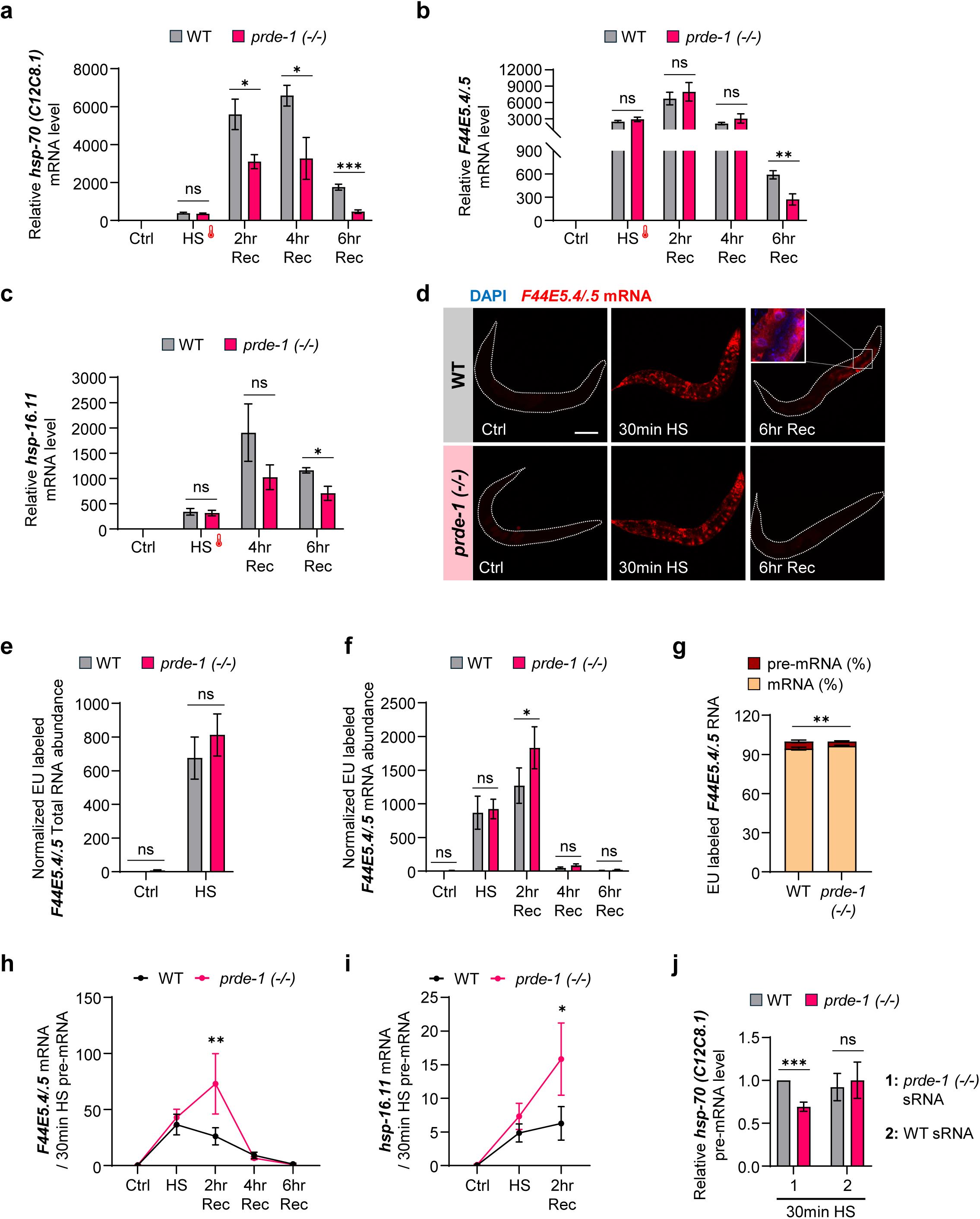
Loss of PRDE-1 accelerates splice completion and alters heat-shock transcript dynamics. **a–c.** RT–qPCR analysis of *hsp-70 (C12C8.1)* **(a)**, *F44E5.4/.5* **(b)**, and *hsp-16.11* **(c)** mRNA in wild-type and *prde-1* animals under control conditions, after 30 min heat shock (HS), and during recovery (Rec). *hsp* mRNAs decline more rapidly during recovery in *prde-1* animals. *n* = 3–13 biological repeats. **d.** RNA FISH of *F44E5.4/.5* mRNA (red) in wild-type and *prde-1* animals under control conditions, after 30 min HS, and following 6 hr recovery. DNA, DAPI (blue). Scale bar, 100 μm. **e–g,** Metabolic EU pulse–chase analysis of *F44E5.4/.5* RNA. **(e)** Total EU-labeled RNA and **(f)** mature EU-labeled mRNA in wild-type and *prde-1* animals during heat shock and recovery. **(g)** Relative proportions of EU-labeled *F44E5.4/.5* pre-mRNA and mature mRNA immediately after 30 min HS. *n* = 5–13 biological repeats. **h, i.** Splice-completion kinetics calculated as the ratio of mature mRNA to pre-mRNA for *F44E5.4/.5* (**h**) and *hsp-16.11* (**i**) in wild-type and *prde-1* animals during heat shock and recovery. Loss of PRDE-1 increases the accumulation of spliced relative to unspliced RNA during recovery. *n* = 5–14 biological repeats. **j.** Relative *hsp-70 (C12C8.1)* pre-mRNA abundance after 30 min HS in wild-type and *prde-1* animals following injection of small-RNA fractions isolated from heat-shocked *prde-1* (1) or wild-type (2) animals. *hsp-70* pre-mRNA levels, normalized to *pmp-3*, relative to heat-shocked wild-type animals injected with small RNAs from *prde-1* mutants. *n* = 6 biological repeats. **a–c, e–j**: Data are mean ± SEM. Statistical analyses: two-tailed unpaired t-test. (**a–c, g, j**); two-way ANOVA (**e, f, h, i**); two-tailed unpaired t-test (**d, f**); Fisher’s exact test (**i**); χ² test (**j**). ns, not significant; ✱ *p* < 0.05; ✱✱ *p* < 0.01; ✱✱✱ *p* < 0.001.

**Supplementary Figure 5.**
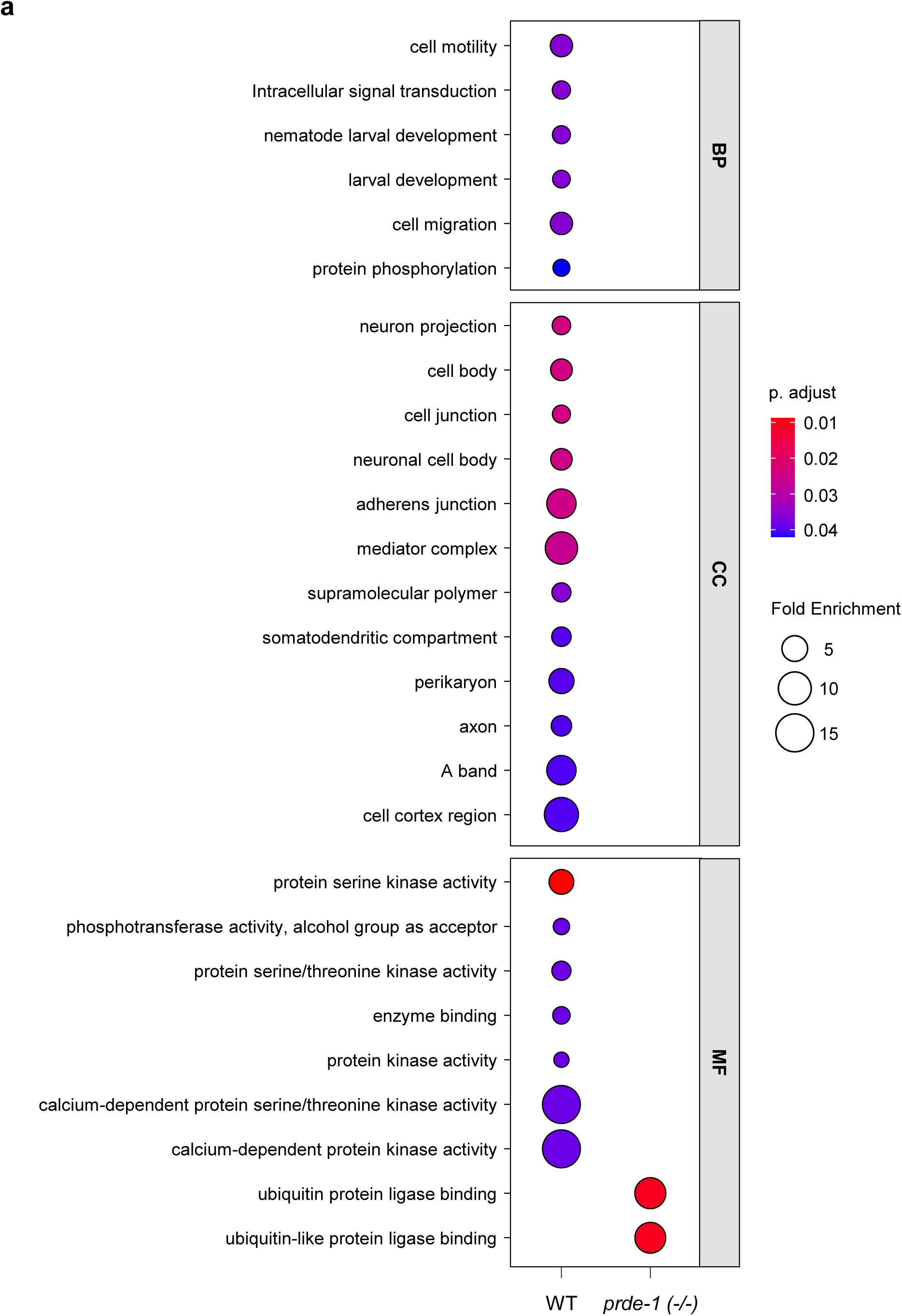
Functional enrichment of heat-shock-regulated alternatively spliced genes targeted by 22G-RNAs. **a,** Gene Ontology (GO) enrichment analysis of genes exhibiting significant alternative splicing that also overlap with 22G-RNA targets in wild-type (WT) and *prde-1* animals. Enriched terms are grouped by biological process (BP), cellular component (CC), and molecular function (MF). Circle size indicates fold enrichment and color indicates Benjamini–Hochberg-adjusted *P* value.

**Supplementary Figure 6.**
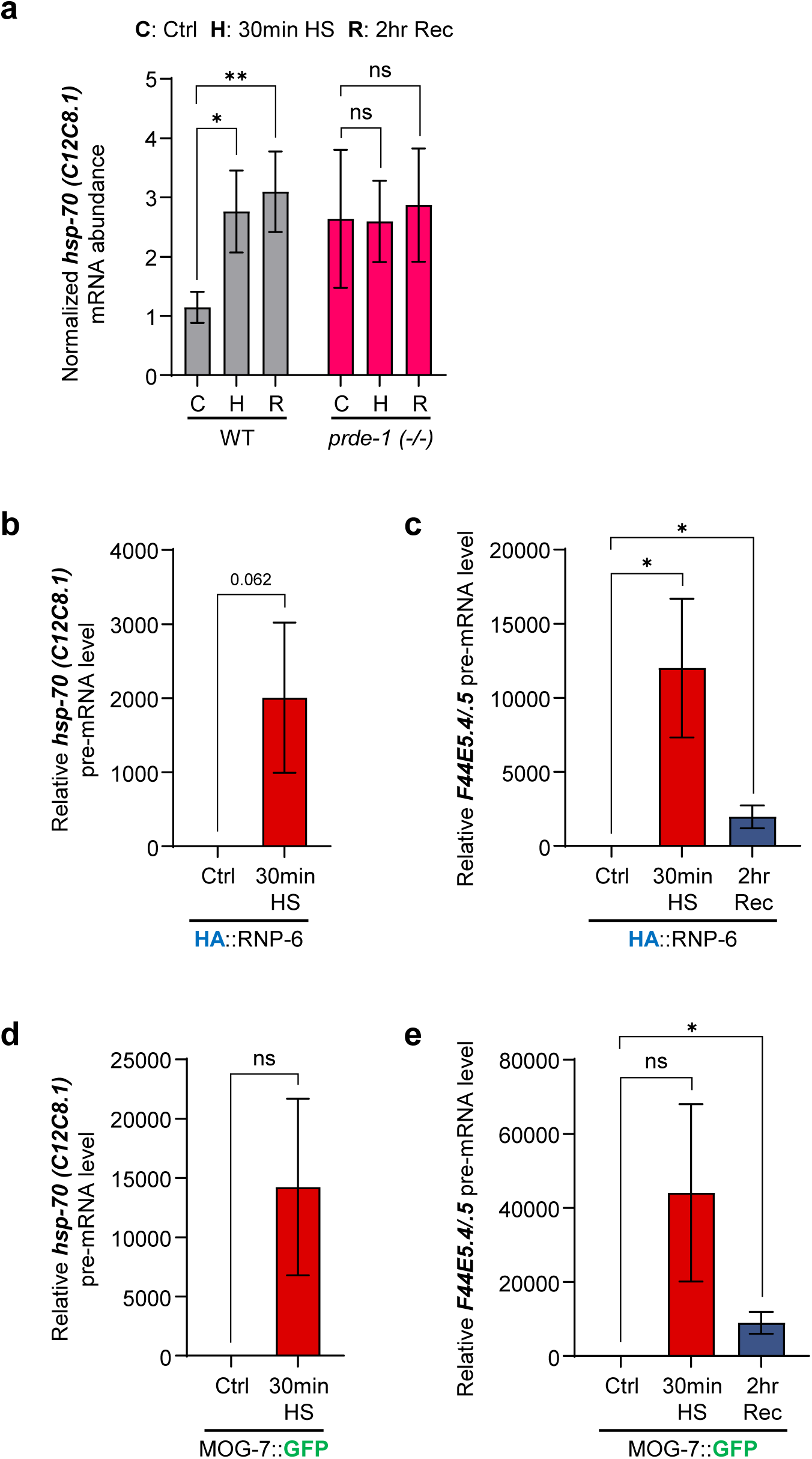
Heat-shock transcripts associate RNA Polymerase II and spliceosomal proteins at the transcription complex. **a.** *hsp* mRNAs co-immunoprecipitated with RNA polymerase II following 4sU labeling and UV crosslinking. Relative abundance of *hsp-70* (*C12C8.1*) mRNA associated with Pol II in wild-type and *prde-1* animals under control conditions (C), after 30 min heat shock (H), and following 2 hr recovery (R). *n* = 5–10 biological repeats. **b, c.** Association of *hsp-70s* pre-mRNA with the early spliceosome factor HA::RNP-6. Co-immunoprecipitated *hsp-70* (*C12C8.1*) **(b)** and *F44E5.4/.5* **(c)** pre-mRNAs were quantified under control conditions, after 30 min HS, and following 2 hr recovery. *n* = 4–5 biological repeats. **d, e.** Association of *hsp-70s* pre-mRNA with the late spliceosome remodeling factor MOG-7::GFP. Co-immunoprecipitated *hsp-70* (*C12C8.1*) **(d)** and *F44E5.4/.5* **(e)** pre-mRNAs were quantified under control conditions, after 30 min HS, and following 2 hr recovery. *n* = 5–10 biological repeats. **a–e**: Data are mean ± SEM. Statistical analyses: two-tailed unpaired t-test (within strain) (**a**); two-tailed unpaired t-test (**b–e**). ns, not significant; ✱ *p* < 0.05; ✱✱ *p* < 0.01.

**Supplementary Figure 7.**
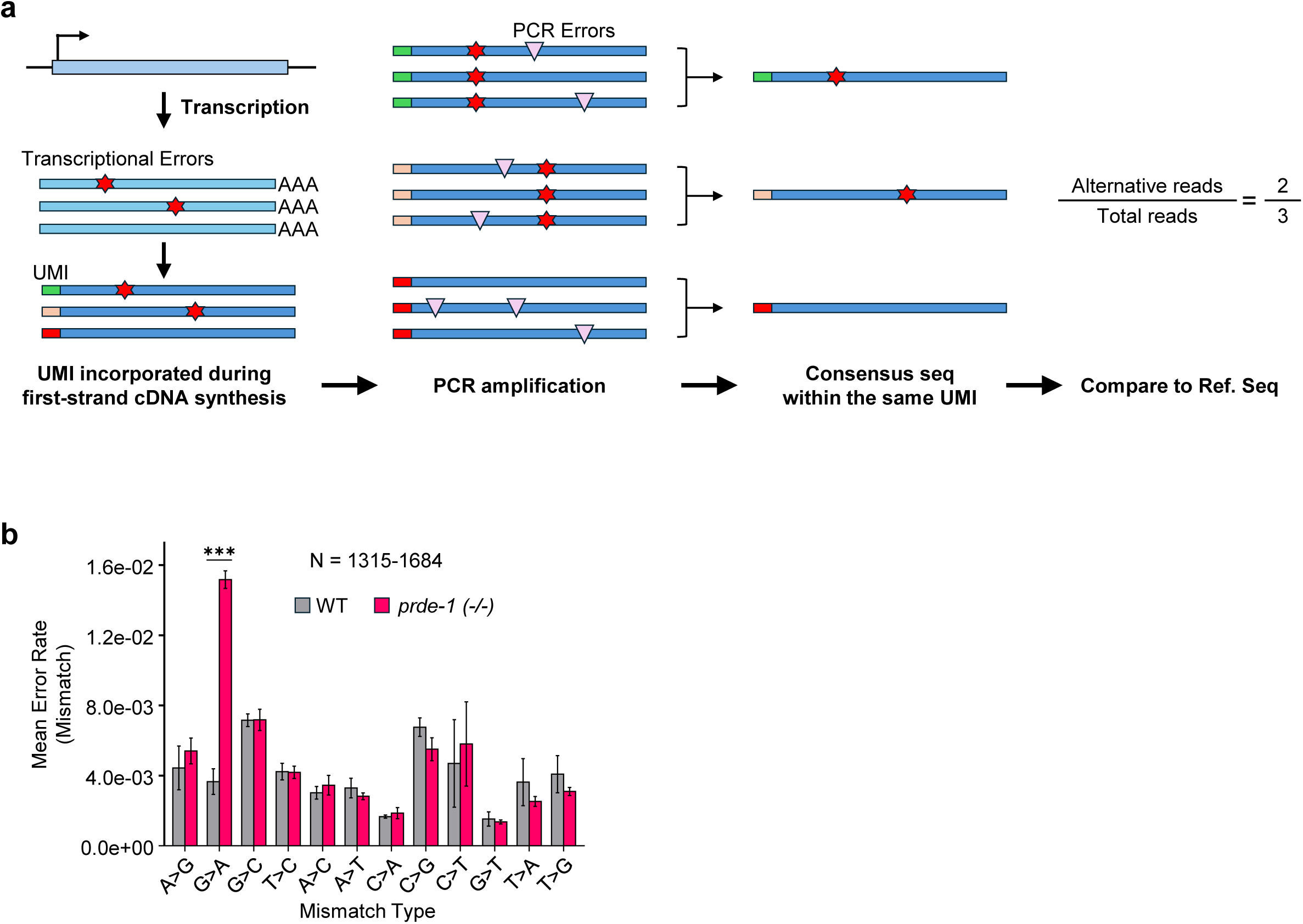
UMI-based transcript error analysis. **a.** Schematic of the unique molecular identifier (UMI)-based approach used to distinguish transcript variants from PCR and sequencing errors. UMIs were incorporated during first-strand cDNA synthesis to uniquely label individual RNA molecules before PCR amplification. Reads sharing the same UMI were collapsed to generate consensus sequences, thereby removing variants introduced during amplification or sequencing. Consensus sequences were compared with the reference sequence to quantify transcript mismatch frequencies. **b.** Frequencies of individual nucleotide substitutions in transcripts from heat-shocked wild-type and *prde-1* animals. Bars show mean mismatch frequency ± SEM across genes grouped by substitution type. G>A substitutions were selectively increased in *prde-1* relative to wild type. *N* = 1,315–1,684 [genes/transcripts—whichever is correct] per analysis. ✱✱✱ *p* < 0.001.

**Supplementary Figure 8.**
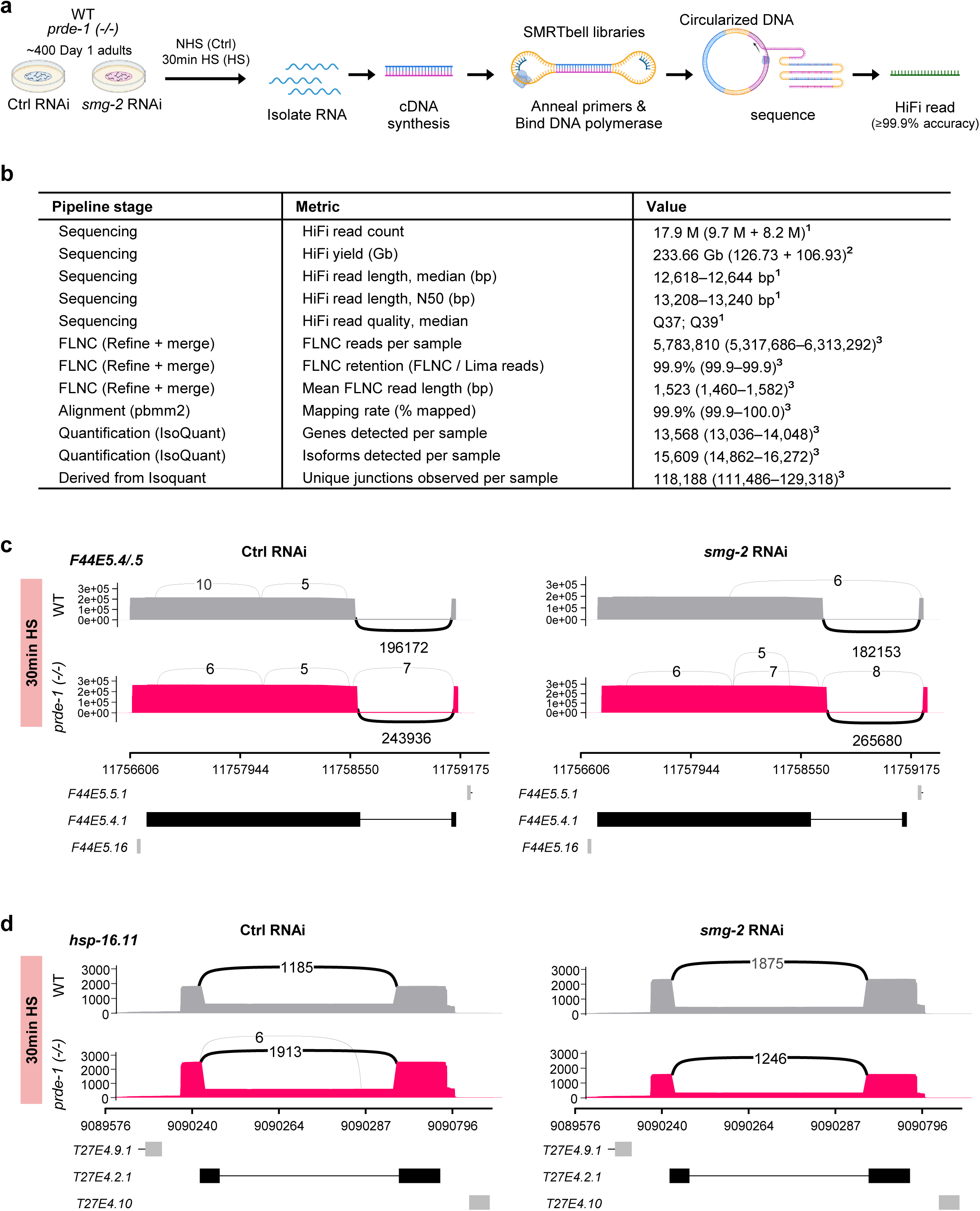
Long-read transcriptome sequencing reveals altered heat-shock transcript processing in *prde-1* mutants. **a.** Experimental design for PacBio HiFi long-read RNA sequencing. Day 1 wild-type and *prde-1* adults were treated with control or *smg-2* RNAi and maintained under control conditions (NHS) or exposed to 30 min heat shock (HS). RNA was converted to cDNA, prepared as SMRTbell libraries, and sequenced to generate high-accuracy HiFi reads. **b.** Summary of long-read sequencing and analysis metrics, including HiFi read yield and quality, full-length non-concatemer (FLNC) read recovery, mapping efficiency, and numbers of genes, isoforms, and unique splice junctions detected. Values summarize sequencing across the biological replicates as indicated. Table notes: **¹** Per SMRT cell., **²** Total across 2 SMRT cells. **³** Median (IQR), *n* = 24. **c, d.** Sashimi plots of long-read coverage and splice-junction usage across *F44E5.4/.5* **(c)** and *hsp-16.11* **(d)** after 30 min HS. Rows show wild-type (gray) and *prde-1* (magenta); columns show control RNAi (left) and *smg-2* RNAi (right). Peaks indicate transcript coverage and arcs indicate splice junctions, with numbers denoting junction-supporting reads. Annotated transcript models are shown below each locus. *prde-1* animals exhibit altered junction usage and greater heterogeneity in transcript structure, including under conditions in which NMD-sensitive transcripts are stabilized by *smg-2* RNAi.

**Supplementary Figure 9.**
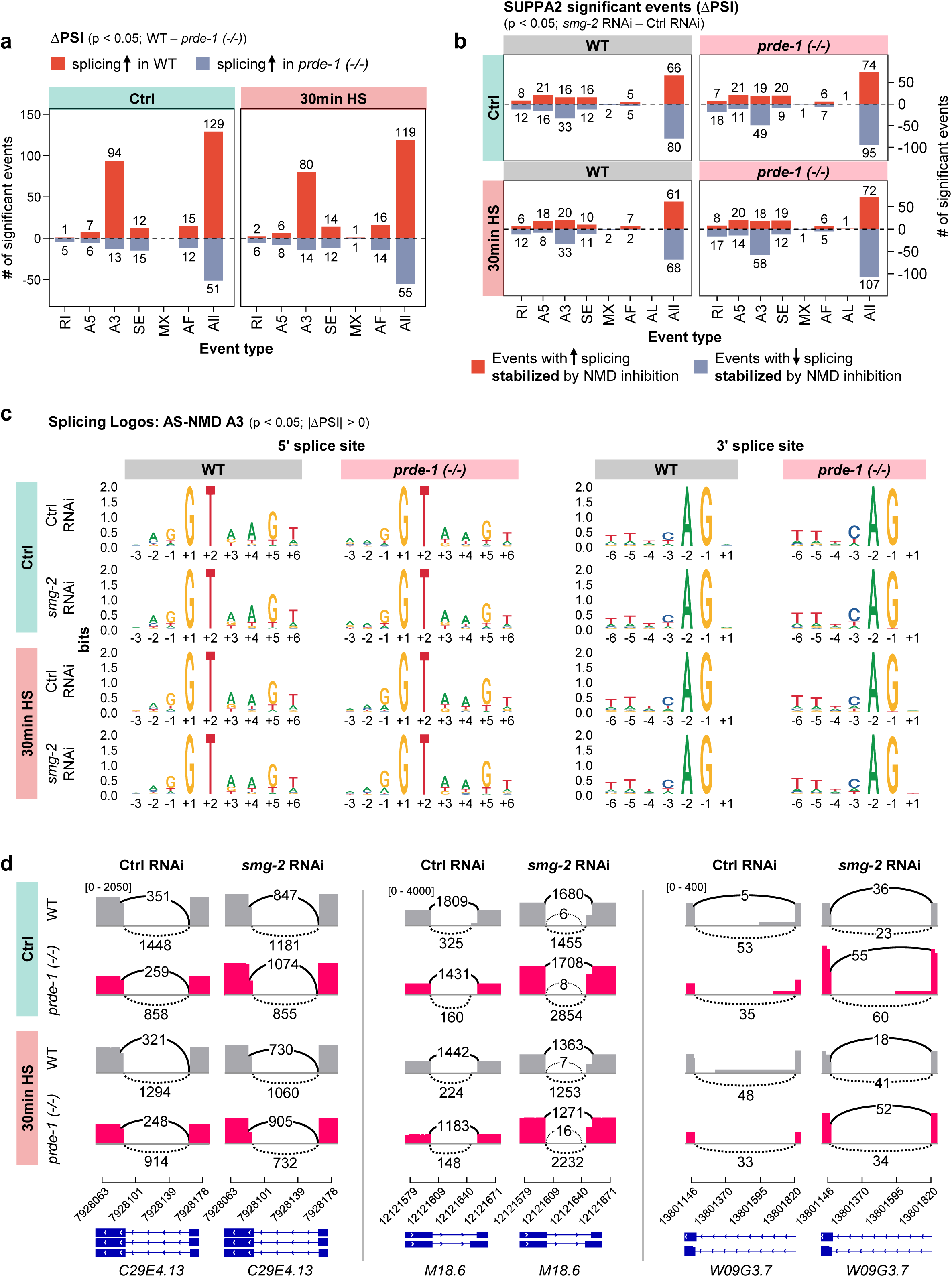
Loss of PRDE-1 alters alternative splice-site selection without disrupting splice-site recognition. **a.** Significant alternative splicing events identified by SUPPA2 in wild-type and *prde-1* animals under control conditions and after 30 min heat shock (HS) with NMD intact. Bars indicate events with increased splicing in wild type (red) or *prde-1* (blue) based on signed ΔPSI (*P* < 0.05). Numbers indicate significant events for each class: retained intron (RI), alternative 5′ or 3′ splice site (A5, A3), skipped exon (SE), mutually exclusive exons (MX), alternative first exon (AF), alternative last exon (AL), and all events combined. **b.** Alternative splicing events stabilized by NMD inhibition. SUPPA2 analysis shows significant AS-NMD events (ÄPSI, *smg-2* RNAi − control RNAi; *P* < 0.05) in wild-type and *prde-1* animals under control conditions (top) and after 30 min HS (bottom). Red and blue bars indicate events with increased or decreased inclusion, respectively, upon NMD inhibition. Numbers indicate significant events in each class. **c.** Sequence logos surrounding the 5′ and 3′ splice sites of NMD-sensitive A3 events in wild-type and *prde-1* animals under control and heat-shock conditions, with or without *smg-2* RNAi. Canonical splice-site consensus sequences are maintained across genotype and treatment, indicating that loss of PRDE-1 alters selection among alternative splice sites rather than splice-site recognition itself. **d.** Representative long-read splice-junction profiles illustrating NMD-sensitive alternative 3′ splice-site usage in *C29E4.13*, *M18.6*, and *W09G3.7*. Junction usage is shown for wild-type (gray) and *prde-1* (magenta) animals under control conditions or after 30 min HS, with control or *smg-2* RNAi. Solid and dashed arcs indicate alternative junctions; numbers denote junction-supporting reads. Gene models are shown below.

**Supplementary Figure 10.**
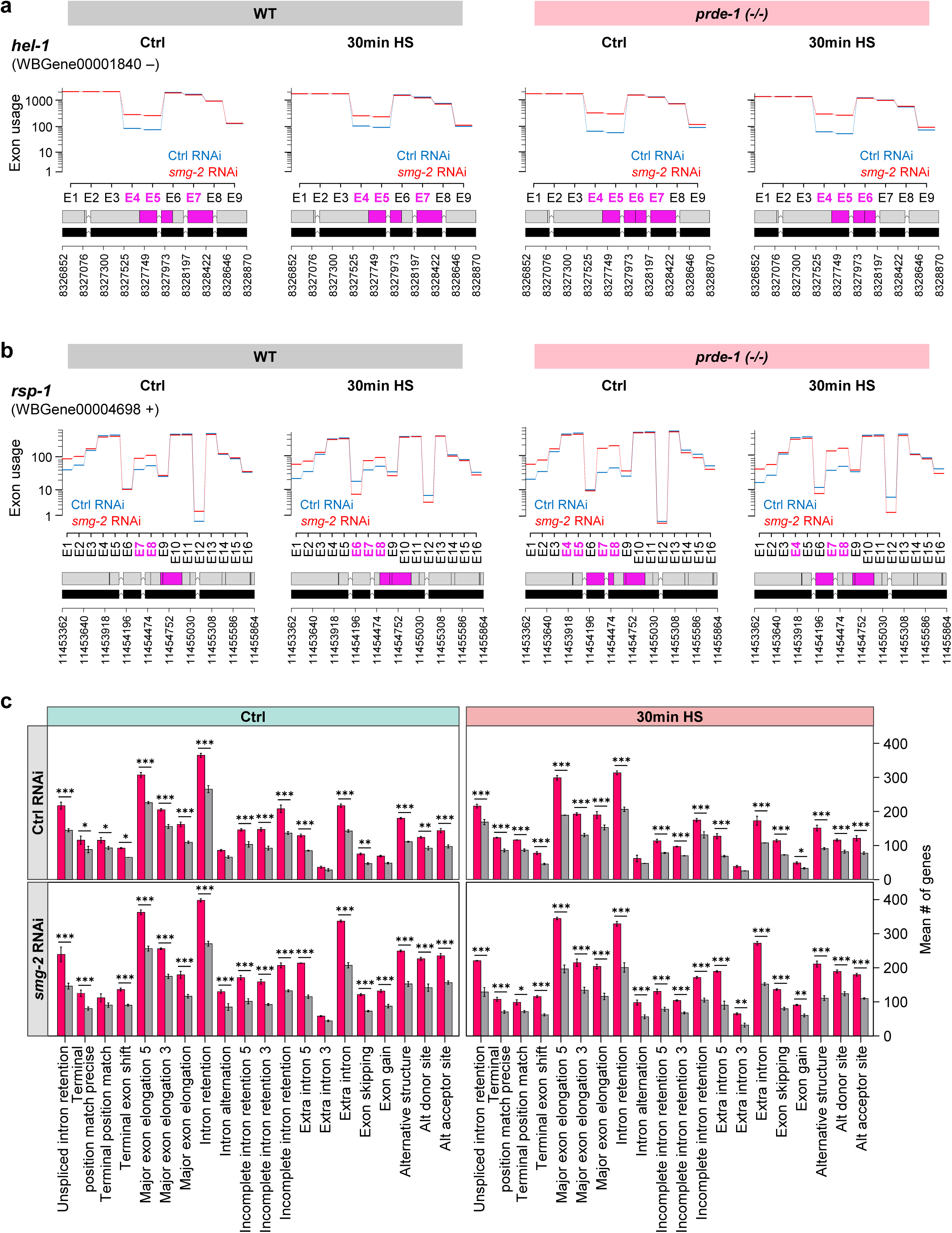
Loss of PRDE-1 increases NMD-sensitive alterations in exon usage and transcript architecture. **a, b.** Differential exon usage in the spliceosomal factors *hel-1* (**a**) and *rsp-1* (**b**) determined by DEXSeq. Exon usage is shown for wild-type and *prde-1* animals under control conditions or after 30 min heat shock (HS), with control RNAi (blue) or *smg-2* RNAi (red). Exons exhibiting significant differential usage upon NMD inhibition are highlighted in magenta. Gene models and genomic coordinates are shown below. **c.** Transcriptome-wide classification of structural changes revealed by NMD inhibition in wild-type (gray) and *prde-1* (magenta) animals under control conditions (left) and after 30 min HS (right). Bars show the number of genes exhibiting each class of transcript alteration under control RNAi (top) or *smg-2* RNAi (bottom), including intron retention, alternative exon usage, alternative splice-site selection, exon skipping, and other changes in transcript structure. Bars represent mean ± SEM. *P* values compare wild type and *prde-1*; ✱ *p* < 0.05; ✱✱ *p* < 0.01; ✱✱✱ *p* < 0.001.

## Notes

### Competing Interest Statement

The authors have declared no competing interest.

### Summary of Updates

Included new data regarding: 1. IP with spliceosome proteins 2. long-read sequencing to show that piRNAs regulate AS-NMD 3. included new experiments showing prg-1 dependence

